# Antibody-drug conjugates targeting CD45 plus Janus kinase inhibitors effectively condition for allogeneic hematopoietic stem cell transplantation

**DOI:** 10.1101/2020.10.02.324475

**Authors:** Stephen P. Persaud, Julie K. Ritchey, Jaebok Choi, Peter G. Ruminski, Matthew L. Cooper, Michael P. Rettig, John F. DiPersio

## Abstract

Despite the curative potential of hematopoietic stem cell transplantation (HSCT), transplant conditioning-associated toxicities preclude broader clinical application. Antibody-drug conjugates (ADC) provide an attractive approach to HSCT conditioning that minimizes toxicity while retaining efficacy. Initial studies of ADC conditioning have largely involved syngeneic HSCT; however, for treatment of acute leukemias or tolerance induction for solid organ transplantation, strategies for allogeneic HSCT (allo-HSCT) are needed. Using murine allo-HSCT models, we show that combining CD45-targeted ADCs with the Janus kinase inhibitor baricitinib enables multilineage alloengraftment with >80-90% donor chimerism. Mechanistically, baricitinib impaired T and NK cell survival, proliferation and effector function, with NK cells being particularly susceptible due to inhibited IL-15 signaling. Unlike irradiated mice, CD45-ADC-conditioned mice did not manifest graft-versus-host alloreactivity when challenged with mismatched T cells. Our studies demonstrate novel allo-HSCT conditioning strategies that exemplify the promise of immunotherapy to improve the safe application of HSCT for treating hematologic diseases.

## INTRODUCTION

Hematopoietic stem cell transplantation (HSCT) has therapeutic potential for hematologic malignancies^1^, autoimmunity^2^, immunodeficiency,^3^ chronic infection^4^, or tolerance induction for solid organ transplantation (SOT)^5^. However, two formidable barriers must be overcome to achieve successful HSCT outcomes. First, recipient-derived hematopoietic stem cells (HSCs) must be depleted to create space for incoming donor HSCs. Second, in allogeneic HSCT (allo-HSCT), host and donor immune responses must be controlled to prevent graft rejection and graft-versus-host-disease (GvHD), respectively^6^. To overcome these barriers, HSCT patients undergo conditioning regimens comprised of chemotherapy and/or irradiation,^7^ whose toxicities limit the use of HSCT to life-threatening conditions like acute myeloid leukemia (AML)^8^.

For AML, allo-HSCT offers the best chance for disease control. Donor T lymphocytes in the HSC allograft mediate graft-versus-leukemia (GvL) effects that protect against relapse^9^. Myeloablative conditioning is preferable for AML as its antileukemia activity also mitigates relapse risk^10^. However, since the median age at diagnosis for AML is 68^11^, patients’ medical comorbidities or functional status may prevent them from undergoing this potentially curative therapy^12^. Moreover, older AML patients have cytogenetically and clinically higher-risk disease that is more treatment-resistant and relapse-prone^13, 14, 15^. This presents a clinical dilemma: the patients most likely to suffer from AML with adverse features are those who most require aggressive therapy, yet they are often the least able to tolerate it.

Novel allo-HSCT conditioning approaches that avoid treatment-related toxicities without sacrificing therapeutic efficacy are urgently needed. Recently, conditioning strategies have emerged using antibody-drug conjugates (ADCs) to target the hematopoietic niche. Initial studies used ADCs comprised of the ribosome inactivator saporin^16^ linked to antibodies recognizing the phosphatase CD45^17^ (CD45-SAP) or the tyrosine kinase c-Kit (cKit-SAP)^18^ to specifically deplete HSCs. In mouse models, CD45-SAP and cKit-SAP were well-tolerated and effectively permitted syngeneic HSCT with high-level donor chimerism. Moreover, these conditioning regimens were used therapeutically in mouse models of sickle cell disease^17^, hemophilia^19^, Fanconi anemia^20^, and recombinase-activating gene (RAG) deficiency^21^.

Fewer studies, however, have studied ADCs as conditioning for allo-HSCT, in which T- and/or NK cell-mediated rejection must be overcome to enable engraftment. Such studies are critical for applying ADC-based conditioning to AML or for tolerance induction in SOT. Prior reports using cKit-targeted regimens have achieved engraftment in major histocompatibility complex (MHC)-mismatched allo-HSCT models^22, 23^. Herein, we used CD45-SAP to develop minimally-toxic conditioning regimens for murine allo-HSCT, with particular emphasis on how these therapies impact host and donor immunity. Using minor histocompatibility antigen (miHA)- and MHC-mismatched models, we demonstrate that CD45-SAP plus pan-T cell depletion (TCD) is sufficient to permit allogeneic donor engraftment. Furthermore, the selective and balanced Janus kinase 1/2 (JAK1/2) inhibitor baricitinib, previously shown to prevent GvHD while enhancing GvL effects^24^, permits robust alloengraftment after CD45-SAP conditioning without requiring pan-TCD. Finally, unlike total body irradiation (TBI) conditioning, CD45-SAP did not promote pathogenic graft-versus-host alloreactivity in mice challenged with allogeneic splenocytes. Taken together, our study provides a novel strategy for allo-HSCT whose biological effects – reducing rejection and GvHD while sparing GvL activity – provide the ideal blend of immunomodulatory activities for the treatment of AML.

## RESULTS

### CD45 and cKit antibody-drug conjugates for syngeneic HSCT conditioning

To evaluate saporin-conjugated CD45 and cKit antibodies as conditioning agents for allo-HSCT, we compared their previously described abilities to deplete murine HSCs and promote syngeneic HSCT^17, 18^. *In vitro*, CD45-SAP and cKit-SAP inhibited hematopoietic colony formation with picomolar-range IC50 values (Figure 1a). Both ADCs effectively depleted HSCs *in vivo*, as defined phenotypically (LSK CD48^-^CD150^+^) or by colony formation (Figure 1b). Importantly, HSC depletion required an intact ADC comprised of the relevant antibody linked to saporin; controls lacking either of these components were devoid of activity. As previously reported, CD45-SAP was strongly lymphodepleting, whereas cKit-SAP lacked this activity (Supplementary Figure 1a). Notably, reduced CD4^+^ and CD8^+^ T cells and increased granulocytes were noted in control mice receiving sAV-SAP or IgG-SAP (Supplementary Figure 1a). These effects were more pronounced in the CD45-SAP group, which received higher doses than the cKit-SAP group, possibly reflecting a dose-dependent effect of sAV-SAP itself. Finally, all ADC-treated mice had CBCs within the reference range (Supplementary Figure 1b).

**Figure 1.**
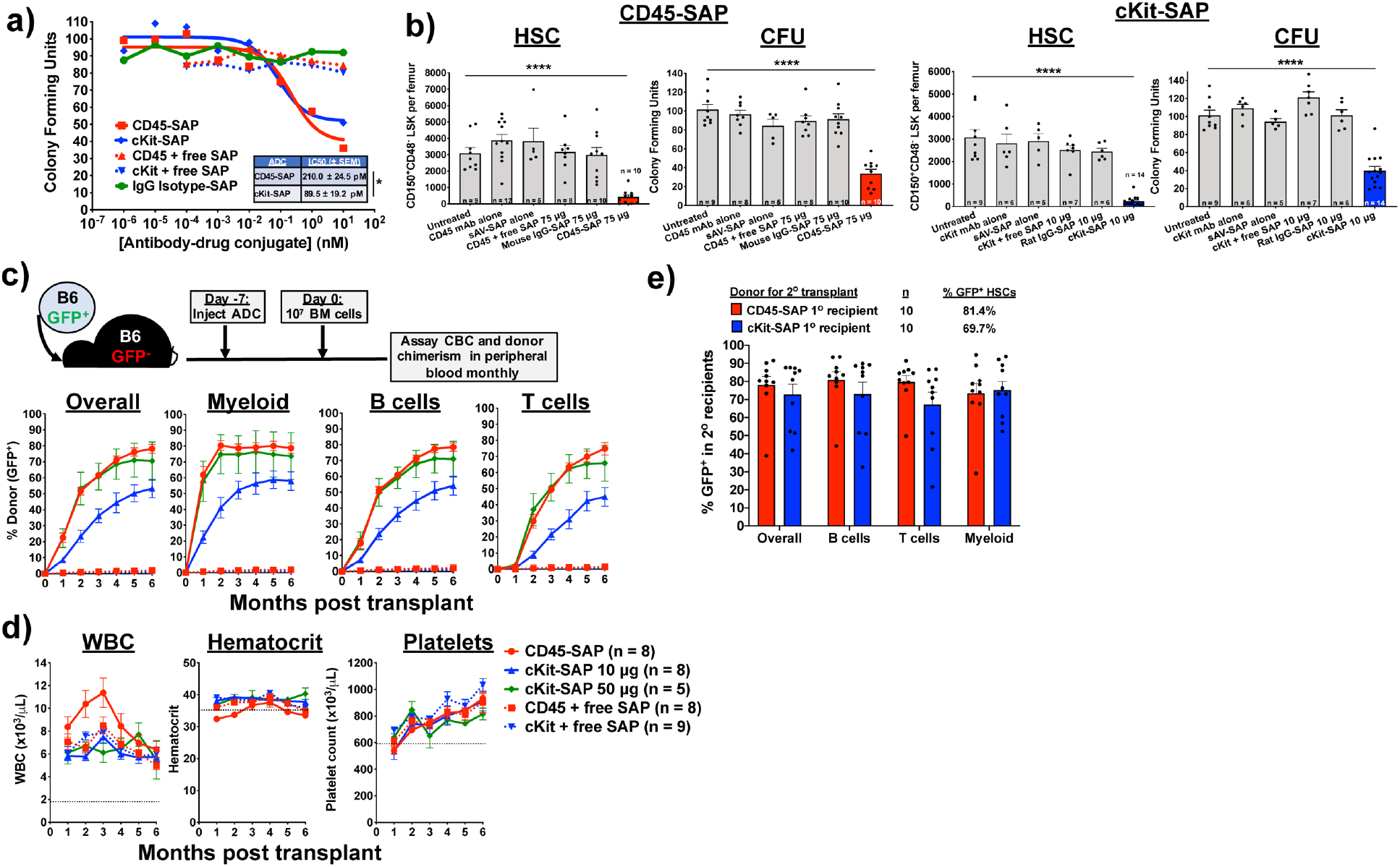
CD45-SAP and cKit-SAP are similarly effective conditioning agents for syngeneic HSCT. **(a)** Inhibition of B6 bone marrow colony formation *in vitro* by ADCs or control conjugates. Mean colony counts from one representative of three experiments are shown. **(b)** *In vivo* depletion of bone marrow CD150^+^CD48^-^ LSK cells (HSC) and colony forming units (CFU) 7 days post-infusion with the indicated conjugates. Mice were pooled from 2-4 experiments; please note that the same cohort of untreated mice was used to compare with the CD45-SAP and cKit-SAP treatment groups. **(c and d)** Schema and results for syngeneic HSCT in mice conditioned with the indicated conjugates. Donor chimerism overall and for T, B, and myeloid (Gr1^+^ and/or CD11b^+^) lineages (c) and CBCs (d) are displayed. Mice were pooled from 2-3 experiments. Overall donor chimerism between active and inactive ADC was significantly different at all timepoints (p < 0.0001 for CD45-SAP vs. CD45 + free SAP; p < 0.001 for cKit-SAP 10 μg vs. cKit + free SAP 10 μg). **(e)** Secondary HSCT using whole marrow from B6-GFP→B6 primary recipients that were conditioned with the indicated ADCs, analyzed at 4 months post-transplant. The %GFP^+^ of HSCs infused to the secondary recipients is shown; mice were pooled from 2 experiments. Data points and error bars represent mean ± SEM. For statistical comparisons, ns = not significant, * = p<0.05, ** = p<0.01, *** = p<0.001, **** = p<0.0001.

In a syngeneic HSCT model (B6-GFP→B6), 3 mg/kg CD45-SAP (75 μg) was well-tolerated and permitted stable, high-level donor engraftment comparable to that reported previously^17^ (Figure 1c). Although cKit-SAP depleted HSCs as effectively as CD45-SAP, even when dosed at 0.4 mg/kg (10 μg), it was somewhat less effective at promoting engraftment. When the cKit-SAP dose was increased to 2 mg/kg (50 μg), donor engraftment of all lineages was equivalent to that seen with CD45-SAP. Donor chimerism in spleen, bone marrow and thymus mirrored that observed in peripheral blood (Supplementary Figure 2a). Finally, successful serial transplantation of bone marrow from CD45-SAP and cKit-SAP conditioned primary transplant recipients confirmed that these primary recipients had engrafted functional HSCs (Figure 1e, Supplementary Figure 2b). Taken together, these studies confirm the efficacy of CD45-SAP and cKit-SAP as conditioning for HSCT in the absence of immunologic barriers.

### CD45-SAP plus *in vivo* T cell depletion enables engraftment in miHA- and MHC-mismatched allo-HSCT

To investigate the efficacy of ADCs for allo-HSCT conditioning, we used two transplant models (Figure 2a): a miHA-mismatched BALB/c-Ly5.1→DBA/2 model and a haploidentical CB6F1→B6 (F1-to-parent) model MHC-mismatched for H-2^d^ in the host-versus-graft direction. We chose CD45-SAP for conditioning in these studies to leverage its lymphodepleting activity to overcome graft rejection. However, CD45-SAP alone was insufficient to allow donor engraftment in either model, suggesting that stronger immunosuppression was necessary.

**Figure 2.**
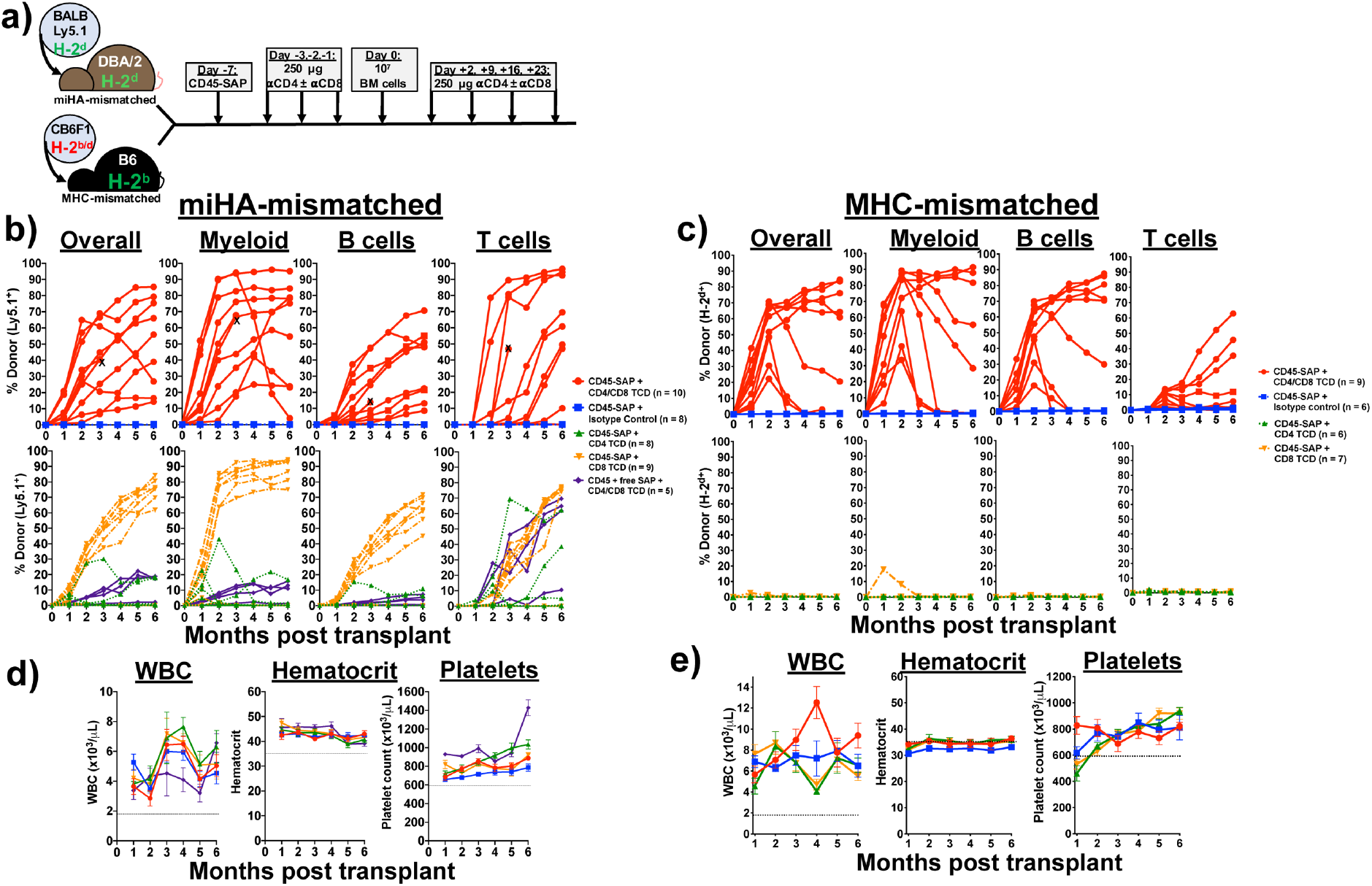
αβ T cell depletion in CD45-SAP conditioned mice permits engraftment in miHA- and MHC-mismatched allo-HSCT. **(a)** Schema for miHA- and MHC-mismatched allo-HSCT models utilizing CD4^+^ and CD8^+^ T cell depletion (TCD) during the peritransplant period. **(b and c)** Peripheral blood donor chimerism for individual mice in the miHA-mismatched (b) and MHC-mismatched alloHSCT models (c). Overall donor chimerism in CD4/CD8 TCD mice was significantly higher than mice receiving isotype control (miHA-mismatched model: p < 0.0001 all timepoints; MHC-mismatched model: p < 0.0001 month 2, p < 0.01 all other timepoints). Data point marked with “X” indicates mouse euthanized for severe head tilt unrelated to the experimental treatment. **(d and e)** Serial CBCs for miHA- (d) and MHC-mismatched (e) models. Data points and error bars in panels (d) and (e) represent mean ± SEM. For statistical comparisons: ns = not significant, * = p<0.05, ** = p<0.01, *** = p<0.001, **** = p<0.0001.

To achieve a fuller, sustained T cell ablation, we treated CD45-SAP conditioned animals with depleting CD4^+^ and/or CD8^+^ antibodies throughout the peritransplant period (Figure 2a)^25^. In the miHA model, while *in vivo* CD4^+^ TCD did not permit significant engraftment, CD8^+^ TCD was sufficient to observe engraftment in 7 of 9 recipient mice. *In vivo* CD4^+^ and CD8^+^ pan-TCD of CD45-SAP conditioned mice resulted in multilineage engraftment in all treated mice, albeit with significant variability in donor chimerism (Figure 2b). Serial CBCs showed stable counts in all lineages (Figure 2d). Gradual loss of donor chimerism was noted in only 1 of 10 pan-TCD mice with myeloid cells declining faster than the longer-lived B cells, a pattern suggestive of inability to engraft or maintain long-term HSCs. Some persistent, low-level donor engraftment, comprised mostly of T cells, was observed in pan-TCD mice conditioned with an inactive ADC. Finally, serial transplantation studies using marrow from CD45-SAP-conditioned, pan-TCD recipients confirmed that these mice engrafted functional donor-derived HSCs (Supplementary Figure 3).

In the MHC-mismatched model, CD4^+^ and CD8^+^ pan-TCD was required for engraftment (Figure 2c). High-level donor chimerism of B cells and myeloid cells and lower T cell chimerism were routinely observed in this system. Although all pan-TCD animals developed donor chimerism in the first two months post-HSCT, 5 of 9 mice showed a gradual loss of donor-derived cells at later timepoints, similar in pace to that seen with one mouse in the miHA model. One recipient showed a sudden, multilineage loss of donor-derived cells, indicative of graft rejection. Serial CBCs were largely stable without any significant periods of post-transplant pancytopenia (Figure 2e).

Although the miHA model has potential for bidirectional alloreactivity, we observed no overt graft rejection or GvHD in miHA-mismatched recipients. This suggested that donor and recipient-derived cells coexisted in stable mixed chimerism, a requirement for tolerance induction for SOT. To directly test for allotolerance, we surgically grafted BALB/c or DBA/2 skin into BALB-DBA mixed chimeric mice (Supplementary Figure S4a). Whereas DBA/2 mice that failed to engraft BALB/c HSCs rejected BALB/c skin by 2 weeks post-implantation^26^, BALB-DBA chimeras were tolerant to BALB/c and DBA/2 skin grafts. As a secondary test, we adoptively transferred CFSE-labeled T cells from BALB-DBA chimeras to new cohorts of BALB/c, DBA/2 or CB6F1 mice (Supplementary Figure S4b). Ninety percent of the transferred T cells were Ly5.1^+^ (donor-derived cells from BALB/c-CD45.1 mice) and did not proliferate when infused into either BALB/c or DBA/2 mice. However, these cells proliferated robustly upon infusion into CB6F1 mice heterozygous for the foreign H-2^b^ haplotype. Taken together, these results verify that our mixed chimeric mice develop cross-tolerance to donor and recipient tissue.

### CD45-SAP combined with the JAK1/JAK2 inhibitor baricitinib promotes multilineage engraftment in allo-HSCT recipients without *in vivo* T cell depletion

Our studies using *in vivo* TCD in miHA- and MHC-mismatched allo-HSCT provide proof-of-principle evidence that ADC-based conditioning regimens can permit engraftment provided that immune barriers are sufficiently suppressed. However, the variability of donor chimerism we observed, the high incidence of graft loss, and the potential risk of opportunistic infections, would limit the clinical utility and translatability of a strategy requiring prolonged TCD. We therefore sought to refine our ADC-based allo-HSCT conditioning regimens with these issues in mind.

Prior work from our laboratory demonstrated that the selective JAK1/2 inhibitor, baricitinib, prevents and even reverses established GvHD, while enhancing GvL effects^24^. The complete prevention of GvHD seen with baricitinib phenocopied that seen in IFNgR-deficient mice treated with aIL-6R, implicating these cytokines’ signaling pathways as important targets of baricitinib effect. Interestingly, mice that received baricitinib showed somewhat improved donor chimerism, although this was in lethally-irradiated mice with donor chimerism already near 100%. However, in a fully-mismatched allo-HSCT model utilizing sublethal irradiation for conditioning (Supplementary Figure 5), IFNgR deficiency in donor and/or recipient cells markedly improved donor chimerism. This result suggested that disabling IFNgR permitted engraftment in the context of reduced-intensity conditioning. We hypothesized that baricitinib, which also blocks IFNγR signaling, may promote engraftment in allo-HSCT when combined with CD45-SAP.

We therefore tested baricitinib in our miHA- and MHC-mismatched allo-HSCT models, first using it in lieu of TCD (Figure 3a). CD45-SAP conditioning plus daily baricitinib administered during the peritransplant period was highly effective in the miHA-mismatched model (Figure 3b). Seven of 10 mice engrafted, all of which showed stable multilineage donor chimerism of ~80% overall. In the MHC-mismatched model, however, daily baricitinib treatment plus CD45-SAP led to donor chimerism in four of seven mice that was stable in only one of those four (Figure 3c). Thus, while daily baricitinib treatment was effective at overcoming immune barriers in miHA-mismatched allo-HSCT, it was ineffective in the MHC-mismatched setting.

**Figure 3.**
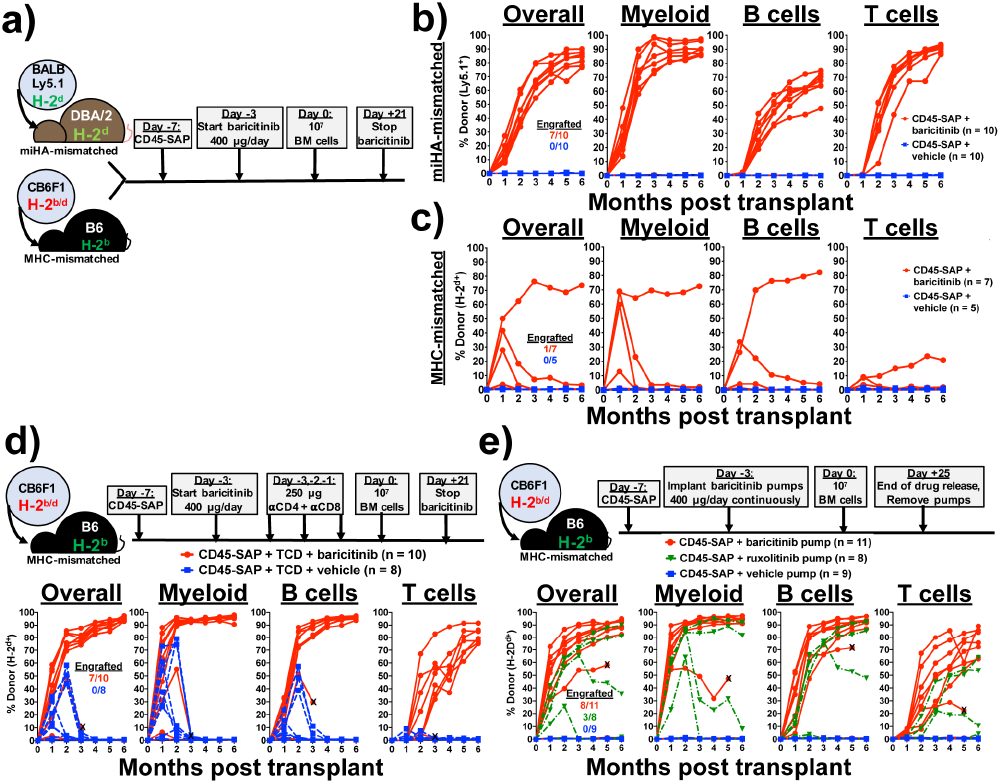
The selective JAK1/2 inhibitor baricitinib permits engraftment in CD45-SAP conditioned mice. **(a)** Schema for baricitinib and CD45-SAP treatment in the miHA- and MHC-mismatched allo-HSCT models. **(b and c)** Peripheral blood donor chimerism for individual mice in the miHA-mismatched (b) and MHC-mismatched (c) models. Differences between baricitinib and vehicle groups were statistically significant at all timepoints in the miHA model (p < 0.001 at month 3, p < 0.01 all other timepoints) and at month 1 only in the MHC-mismatched model (p < 0.01). **(d)** Schema and results for MHC-mismatched HSCT combining CD45-SAP, daily baricitinib, and pre-transplant TCD. Differences between baricitinib and vehicle groups were statistically significant at all timepoints (p < 0.05 months 1-2, p < 0.01 months 3-6). Data point marked with “X” indicates mouse euthanized early to assess rapid loss of donor engraftment. **(e)** Schema and results for MHC-mismatched HSCT combining CD45-SAP conditioning with continuously-infused JAK1/2 inhibitors. Differences between baricitinib and vehicle groups were significant at all timepoints (p < 0.001 months 1-4; p < 0.01 months 5-6); differences between ruxolitinib and vehicle groups were significant at months 1 and 2 (p < 0.0001 and p < 0.001, respectively). Data point marked with “X” indicates mouse death one week prior to collection of final timepoint. Insets represent the numbers of successfully engrafted mice at t = 6 months. For statistical comparisons: ns = not significant, * = p<0.05, ** = p<0.01, *** = p<0.001, **** = p<0.0001.

Since T cells comprise the major barrier to engraftment in the both miHA- and MHC mismatched models, failed engraftment in mice receiving baricitinib daily likely reflects insufficient host T cell immunosuppression to prevent rejection. We reasoned that reducing the strength of the alloresponse by other means may improve baricitinib efficacy. We therefore combined CD45-SAP and daily baricitinib therapy with pre-transplant pan-TCD, essentially substituting in baricitinib for post-transplant TCD (Figure 3d). This regimen was highly effective, achieving stable engraftment in 7 of 10 mice, with overall donor chimerism >90%. The donor chimerism in all lineages, particularly T cells, was superior to that seen with baricitinib or pan-TCD alone. By contrast, mice receiving vehicle instead of baricitinib engrafted temporarily but experienced graft failure or rejection by three months post-HSCT.

Pharmacokinetics may also have impacted the efficacy of baricitinib monotherapy in MHC-mismatched HSCT. Data from a prior study showed that subcutaneous baricitinib has a plasma half-life in B6 mice of approximately one hour^27^, suggesting a prolonged absence of circulating drug if dosed every 24 hours. To test the duration of baricitinib effect, we conducted a pharmacodynamic study in which mice received a single baricitinib dose, then were followed over time with a whole blood assay for IFNγ-induced Stat1 phosphorylation (Supplementary Figure 6). Baricitinib at 400 μg completely suppressed Stat1 phosphorylation at 4 hours post-infusion, an effect that was diminished slightly at 12 and 24 hours and absent by 36 hours. By comparison, 80 μg baricitinib provided only partial suppression at 4 hours post-infusion that was absent at later timepoints.

We hypothesized that a continuous presence of baricitinib would provide more sustained immunosuppression. We therefore administered the same 400 μg daily dose of baricitinib continuously via subcutaneous osmotic pumps. Baricitinib was readily soluble in DMSO mixed 1:1 with PEG-400 (Supplementary Figure 7a), and remained soluble and bioactive in this vehicle after a 30-day incubation at 37°C (Supplementary Figure 7b). Peripheral blood leukocytes from B6 mice implanted with baricitinib-loaded pumps showed impaired Stat1 phosphorylation in response to IFNγ, confirming drug release *in vivo* (Supplementary Figure 7c).

Continuously-infused baricitinib (Figure 3e) was more effective than daily baricitinib (Figure 3c) in promoting multilineage engraftment in MHC-mismatched allo-HSCT, achieving >80% overall donor chimerism in 8 of 11 mice. This appeared to be a class effect of JAK1/2 inhibitors, as the related inhibitor ruxolitinib also permitted engraftment, albeit less effectively than baricitinib. As we consistently observed in the MHC-mismatched model when JAK inhibitors were dosed daily, mice with baricitinib pumps developed a mild anemia at the earliest time points which corrected by the later timepoints; otherwise, CBCs were at or above the lower reference limit (Supplementary Figure 8).

Taken together, these studies demonstrate multiple effective, feasible strategies using CD45-SAP and JAK1/2 inhibitors to achieve high-level donor chimerism in both miHA- and MHC-mismatched allo-HSCT without prolonged, global T cell ablation.

### Baricitinib promotes engraftment via suppression of T and NK cell-mediated rejection

We next pursued the mechanisms by which baricitinib promotes engraftment in allo-HSCT. While we hypothesize that suppression of T cell alloreactivity is an important component, disruption of JAK1/2 signaling may impact engraftment in other ways, such as direct effects on donor hematopoiesis^28, 29, 30^. To investigate the degree to which immunosuppression versus other mechanisms contributes to engraftment, we applied baricitinib to CD45-SAP-conditioned syngeneic HSCT, in which immune barriers to engraftment are absent. In peripheral blood and lymphoid organs (Figures 4a and 4b), no significant difference in donor chimerism was observed between CD45-SAP-conditioned mice receiving baricitinib versus vehicle. Importantly, no engraftment was observed in baricitinib-treated mice conditioned with inactive ADC, indicating that baricitinib alone cannot make space for donor HSCs.

**Figure 4.**
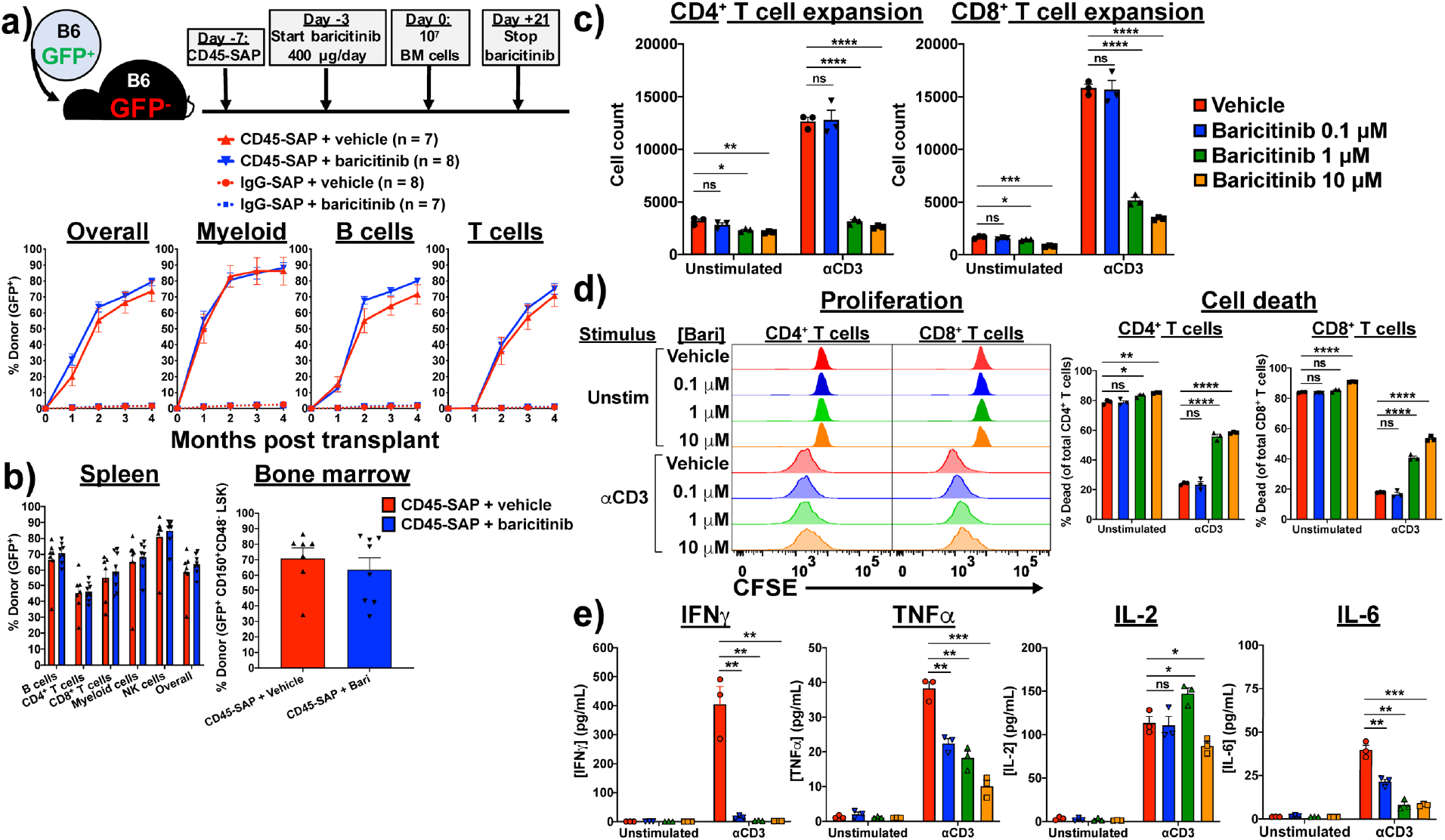
Baricitinib suppresses T cell function and viability, and minimally impacts syngeneic HSCT. **(a)** Schema and results for syngeneic HSCT model in which recipients were conditioned with CD45-SAP or inactive ADC with or without daily baricitinib injections. **(b)** Donor chimerism in spleen and bone marrow of mice from panel (a). **(c)** *In vitro* expansion of αCD3-stimulated (1 μg/mL, 72 hours), CFSE-labeled B6 T cells in the presence of varying concentrations of baricitinib. **(d)** Proliferation and viability of cultures in panel (c). **(E)** Cytokines present in supernatants collected from cultures described in panel (c) after 24 hours incubation. For panels (c-e), data from three technical replicates are shown from one representative of four experiments. For all panels, data points and error bars represent mean ± SEM. For statistical comparisons: ns = not significant, * = p<0.05, ** = p<0.01, *** = p<0.001, **** = p<0.0001.

To characterize the acute effects of baricitinib treatment on HSCT recipients, we analyzed peripheral blood and lymphoid organs of B6 mice that received daily baricitinib for 4 days, the same time period baricitinib is administered before HSCT. Baricitinib treatment minimally affected CBCs or bone marrow cellularity but was associated with a significant reduction in spleen cellularity (Supplementary Figures 9a and 9b). Bone marrow hematopoietic stem and progenitor cell (HSPCs) numbers were largely unaffected by baricitinib, except for somewhat lower frequencies of long-term HSC (CD34^-^CD135^-^ LSK) and megakaryocyte-erythroid progenitors (CD16/32^-^CD34^-^ LK; Supplementary Figure 9c). Absolute myeloid, conventional T cell, and FoxP3^+^ Treg counts were similar in all examined organs, but lower frequencies of B cells were noted in baricitinib-treated mouse spleens (Supplementary Figure 9d and 9e). Finally, immunophenotyping of the splenic T cell and antigen presenting cell (APC) compartments revealed no differences between baricitinib- and vehicle-treated mice (Supplementary Figures 9f and 9g).

To examine how baricitinib impacts T cell responses, we cultured polyclonally-stimulated, CFSE-labeled B6 T cells *in vitro* with baricitinib or vehicle. Baricitinib impaired expansion of αCD3-stimulated CD4^+^ and CD8^+^ T cells in a dose-dependent manner (Figure 4c), due to increased cell death and mildly reduced cell proliferation (Figure 4d). As expected with primary murine cells, unstimulated cultures showed significant T cell death after 72 hours; importantly, the degree of cell death in these cultures was only subtly increased by baricitinib at the highest tested dose, suggesting against nonspecific toxicity. Concentrations of TNFa, IL-6 and, particularly, IFNγ in the culture supernatants were reduced by baricitinib during the culture period (Figure 4e). This reduction was not generalizable, as IL-2 secretion was unaffected by baricitinib. In summary, although baricitinib minimally affects resting T cells in pre-HSCT recipients, activated T cell function is more adversely affected. This is consistent with our hypothesis that baricitinib acts predominantly via immunosuppression, exerting its major therapeutic function on alloreactive T cells that become activated in response to donor HSCs.

In order to extend baricitinib-based conditioning to fully haploidentical (F1→F1) and fully MHC-mismatched models (i.e., BALB/c→B6), inhibition of both T and NK cells is necessary. Multiple reports have shown that ruxolitinib depletes NK cells in mice and humans, impairing NK cell proliferation, cytotoxicity and cytokine production^31, 32, 33^. We hypothesized that baricitinib, via inhibition of JAK1/2 signaling, would show similar biological effects and protect against NK cell-mediated rejection. To test this, we administered CD45-SAP plus baricitinib as conditioning for B6→CB6F1 (parent-to-F1) allo-HSCT (Figure 5a). In this model, engraftment of parental HSCs is resisted by CB6F1 NK cells, which react to the absence of H-2^d^ on the donor-derived cells (“missing self’ recognition)^34^. This phenomenon, termed hybrid resistance, provides an opportunity to isolate NK cell-mediated host-versus-graft responses and investigate how baricitinib affects them.

**Figure 5.**
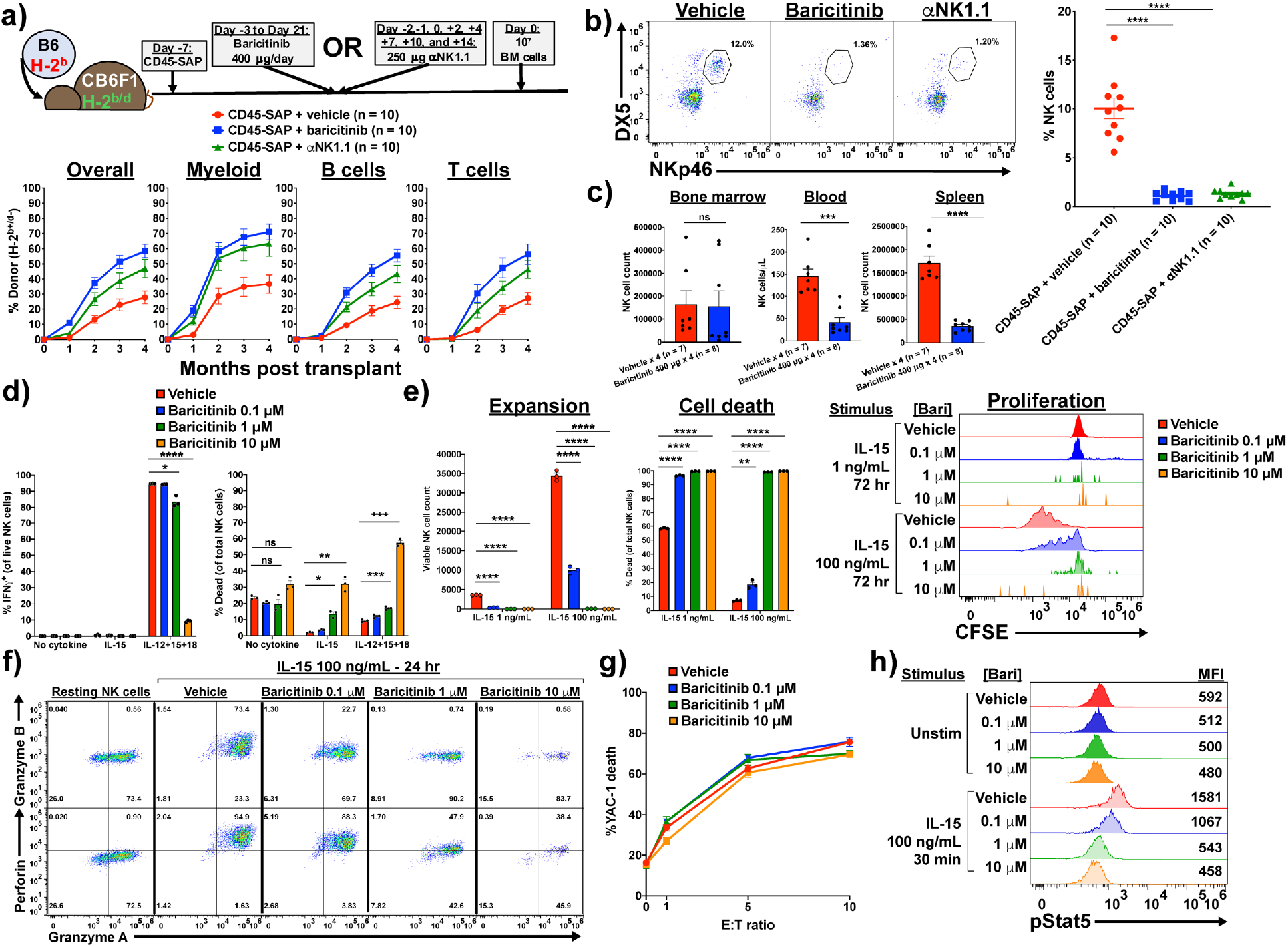
Baricitinib overcomes NK cell-mediated rejection by impairing NK cell survival and effector function. **(a)** Schema and results for parent-to-F1 HSCT model to study baricitinib effect on NK-mediated rejection. Overall peripheral blood donor chimerism was significantly higher for the baricitinib and αNK1.1 groups compared to vehicle at all timepoints (baricitinib vs. vehicle: p < 0.001 months 1, 3 and 4, p < 0.0001 month 2; αNK1.1 vs. vehicle: p < 0.05 months 1-3, p < 0.01 month 4). **(b)** Peripheral blood NK cell frequencies of recipients in panel (a) immediately before HSCT. **(c)** NK cell counts by organ in B6 mice receiving four once-daily doses baricitinib or vehicle. **(d-f)** Functional assays of IL-15-stimulated B6 splenic NK cells incubated with baricitinib or vehicle: IFNg production and survival after 15 hours (d), expansion after 72 hours (e), and cytolytic enzyme expression after 24 hours (f). (**g**) YAC-1 killing by NK cells primed with IL-15 for 48 hours without baricitinib, then washed and plated with target cells for 4 hours with baricitinib or vehicle. **(h)** Stat5 phosphorylation in NK cells after IL-15 stimulation with baricitinib or vehicle present. For panels (d-h), two (h) or three (d-g) technical replicates from one of three experiments are shown; for panel (f), inset numbers are the percentage of events in each quadrant. Data points and error bars represent mean ± SEM. For statistical comparisons: ns = not significant, * = p< 0.05, ** = p<0.01, *** = p<0.001, and **** = p<0.0001.

Overall donor chimerism in B6→CB6F1 transplants treated with CD45-SAP plus vehicle was approximately 25% four months post-HSCT (Figure 5a), considerably lower than that obtained in syngeneic HSCT (Figure 1c). As expected, engraftment was improved by αNK1.1 depletion. Mice conditioned with baricitinib showed overall donor chimerism approaching 60%, surpassing that obtained with αNK1.1 depletion. Pre-HSCT analysis of peripheral blood revealed that both αNK1.1 treatment and baricitinib markedly depleted CB6F1 recipients’ circulating NK cells (Figure 5b). We confirmed this finding in the spleen and peripheral blood, but not bone marrow, of B6 mice treated with daily baricitinib (Figure 5c). Thus, baricitinib overcame NK cell-mediated barriers to HSCT due to efficient *in vivo* NK cell depletion.

NK cell development, maturation, and function depend upon IL-15, which signals through JAK1 and JAK3 to activate Stat5^35^. We asked whether baricitinib disrupts this critical signaling pathway to compromise NK cell survival and function. Murine NK cells stimulated *in vitro* with IL-15 alone or a cocktail of IL-12, IL-15 and IL-18^36^ showed dose-dependent increases in cell death and decreases in IFNg production in response to baricitinib (Figure 5d). As with T cells, nonspecific toxicity in unstimulated cultures was modest and noted only at the highest baricitinib doses. In longer cultures, baricitinib impaired IL-15-mediated NK cell expansion, an effect attributable to dramatically reduced proliferation and viability (Figure 5e). Baricitinib also strongly suppressed IL-15-induced upregulation of the lytic granule enzymes perforin and granzyme B (Figure 5f), which are required for full NK cell cytotoxicity^37^. However, baricitinib did not prevent killing of YAC-1 target cells when added to NK cells that had been already primed with IL-15, suggesting that baricitinib inhibits the acquisition but not the execution of NK cytotoxicity (Figure 5g). Finally, analysis of IL-15 signaling confirmed that baricitinib inhibits IL-15-induced Stat5 phosphorylation in a dose-dependent manner (Figure 5h). These data collectively demonstrate that baricitinib potently impairs NK cell viability, proliferation, and effector function via interference with the IL-15-Stat5 signaling axis.

### CD45-SAP conditioning poorly stimulates pathogenic graft-versus-host alloreactivity compared to TBI

The contribution of conditioning regimen intensity to the development of acute and chronic GvHD is well-studied^38, 39, 40, 41^. A multitude of variables modulate GvHD risk, including donor and recipient age, GvHD prophylaxis, donor HSC source and relatedness, and degree of HLA disparity, which can influence the choice of conditioning intensity^42^. Theoretically, host tissue injury caused by chemotherapy and radiation amplifies GvHD via release of endogenous damage- and pathogen-associated molecular patterns from dying cells. These mediators activate innate immunity, arming APCs to prime vigorous alloreactive T cell responses^43, 44, 45^. We asked whether or not CD45-SAP, with its minimal tissue toxicities, would behave similarly.

To study the effect of conditioning regimen on T cell alloresponses *in vivo*, we used a parent-to-F1 adoptive transfer model, in which alloreactivity is exclusively in the graft-versus-host direction (Figure 6a). In this system, F1 mice conditioned with sublethal irradiation that receive parental splenocytes develop pancytopenia secondary to T cell-mediated marrow aplasia^46^. We compared CD45-SAP to sublethal rather than lethal irradiation (as is typically used in standard GvHD models) to more closely match the severity and degree of myeloablation of the conditioning regimens. Both CD45-SAP and 500 cGy irradiation are nonlethal and have been shown to permit similar levels of syngeneic HSC engraftment^17^, suggesting a similar capacity to generate marrow HSC niche space.

**Figure 6.**
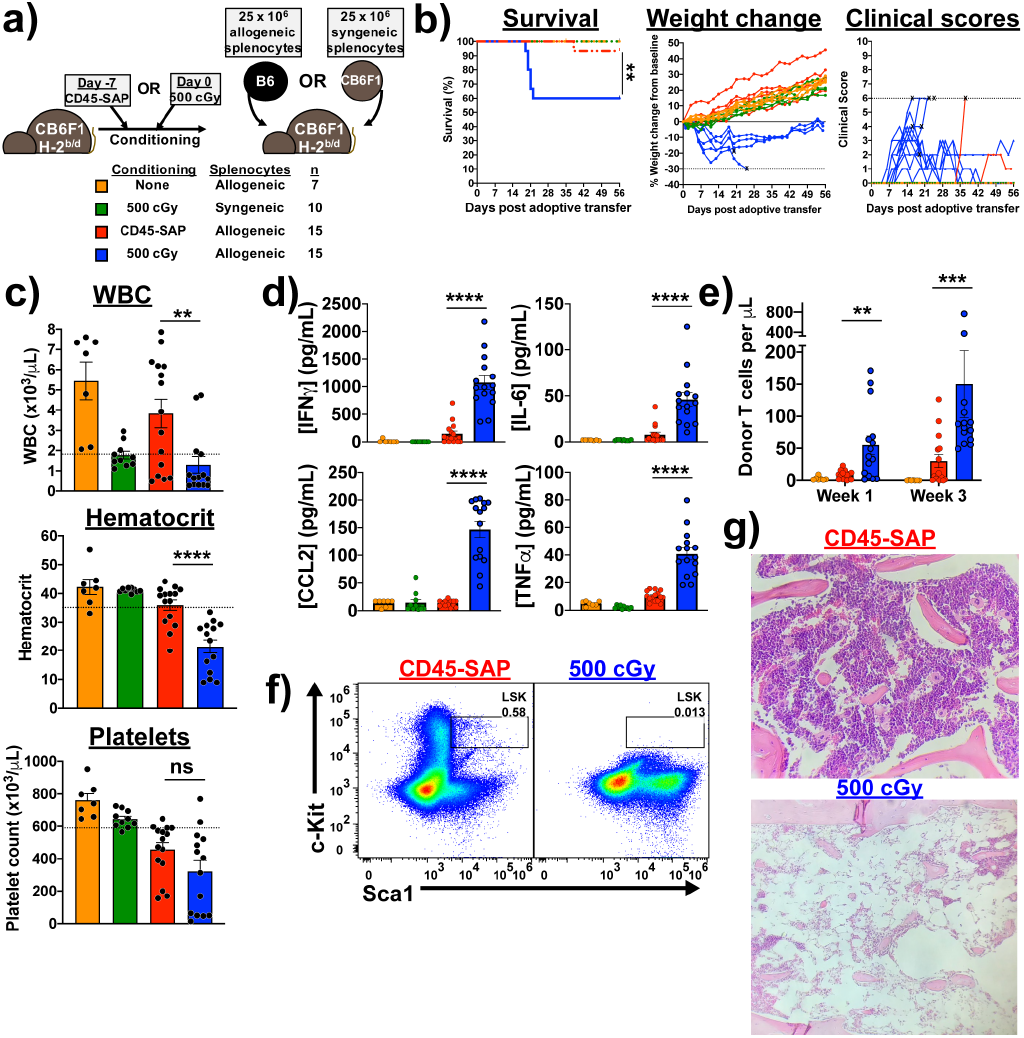
CD45-SAP conditioning does not promote graft-versus-host alloreactivity. **(a)** Schema for parent-to-F1 adoptive transfer model, with sublethal irradiation or CD45-SAP conditioning administered with the usual timing with respect to HSCT. Treatment groups are color-coded throughout the figure per the indicated legend. **(b)** Clinical outcomes for mice treated as per panel (a); “X” indicates death or euthanasia and dotted lines indicate euthanasia thresholds. **(c)** CBCs at 21 days post-splenocyte infusion. **(d)** Plasma inflammatory cytokine concentrations 7 days post-splenocyte infusion. (**e**) Circulating donor T cells at days 7 and 21 post-splenocyte infusion. **(f and g)** Flow cytometry (f; gated on 7-AAD^-^Lineage^-^ cells) and histology (g) of bone marrow from a CD45-SAP conditioned mouse 56 days after allogeneic splenocyte infusion compared with an irradiated mouse that succumbed at day 21. For clarity, weight changes shown in (b) are from a representative sample of five mice per group; for the other plots, all mice analyzed over 2 or 3 independent experiments are included. Data points and error bars represent mean ± SEM. For statistical comparisons: ns = not significant, * = p<0.05, ** = p<0.01, *** = p<0.001, **** = p<0.0001.

Compared to ADC-conditioned mice, TBI-conditioned mice infused with allogeneic splenocytes showed poorer survival and clinical courses along with greater weight loss (Figure 6b). At three weeks post-splenocyte infusion, TBI-conditioned, but not ADC-conditioned mice, developed pancytopenia (Figure 6c) and marked elevations in the plasma concentrations of several inflammatory cytokines, particularly IFNg (Figure 6d). Importantly, TBI-conditioned mice receiving syngeneic splenocytes and unconditioned mice receiving allogeneic splenocytes showed no morbidity, mortality, cytopenias, or pro-inflammatory cytokinemia, confirming that irradiation plus allogeneic T cells are required for pathology. Circulating donor-derived T cells were present in ADC-conditioned mice but at lower frequencies than irradiated mice, indicating that the lack of disease in ADC-conditioned mice is not due to failure of these cells to engraft (Figure 6e). Finally, bone marrow histopathology and flow cytometry demonstrated profound marrow aplasia and HSPC depletion in TBI-conditioned mice that developed lethal disease (Figure 6f and 6g).

To understand why high doses of allogeneic T cells failed to elicit disease in CD45-SAP-conditioned mice, we analyzed the donor T cell response in the lymphoid organs of ADC-versus TBI-conditioned mice. While donor CD4^+^ and CD8^+^ T cells were identified in the spleens of both ADC- and TBI-conditioned mice, the bone marrows of ADC-conditioned mice were virtually devoid of donor T cells (Figure 7a). This contrasted starkly with irradiated mice, whose marrows were extensively infiltrated by donor T cells, mainly CD8^+^ T cells. While higher expression of the bone marrow homing chemokine receptor CXCR4 on CD8^+^ T cells could explain why these were the predominant marrow-infiltrating cells in irradiated mice, differences in CXCR4 expression cannot account for the differences observed between ADC- and TBI-conditioned mice (Figure 7b). Most T cells in ADC- and TBI-conditioned mice had a CD44^hi^CD62L^lo^ effector phenotype, with somewhat higher frequencies in irradiated mice. Donor CD8^+^ T cells in irradiated mice upregulated perforin and granzymes A and B relative to ADC-conditioned mice, indicating greater potential for cytotoxicity (Figure 7c). Higher expression of MHC and the costimulatory receptors CD80 and CD86 (Figure 7d) were noted in host-derived APCs from irradiated mice compared to ADC-conditioned mice. Collectively, these data suggest that ADC-conditioning produces a suboptimal environment for priming a pathogenic allogeneic T cell response.

**Figure 7.**
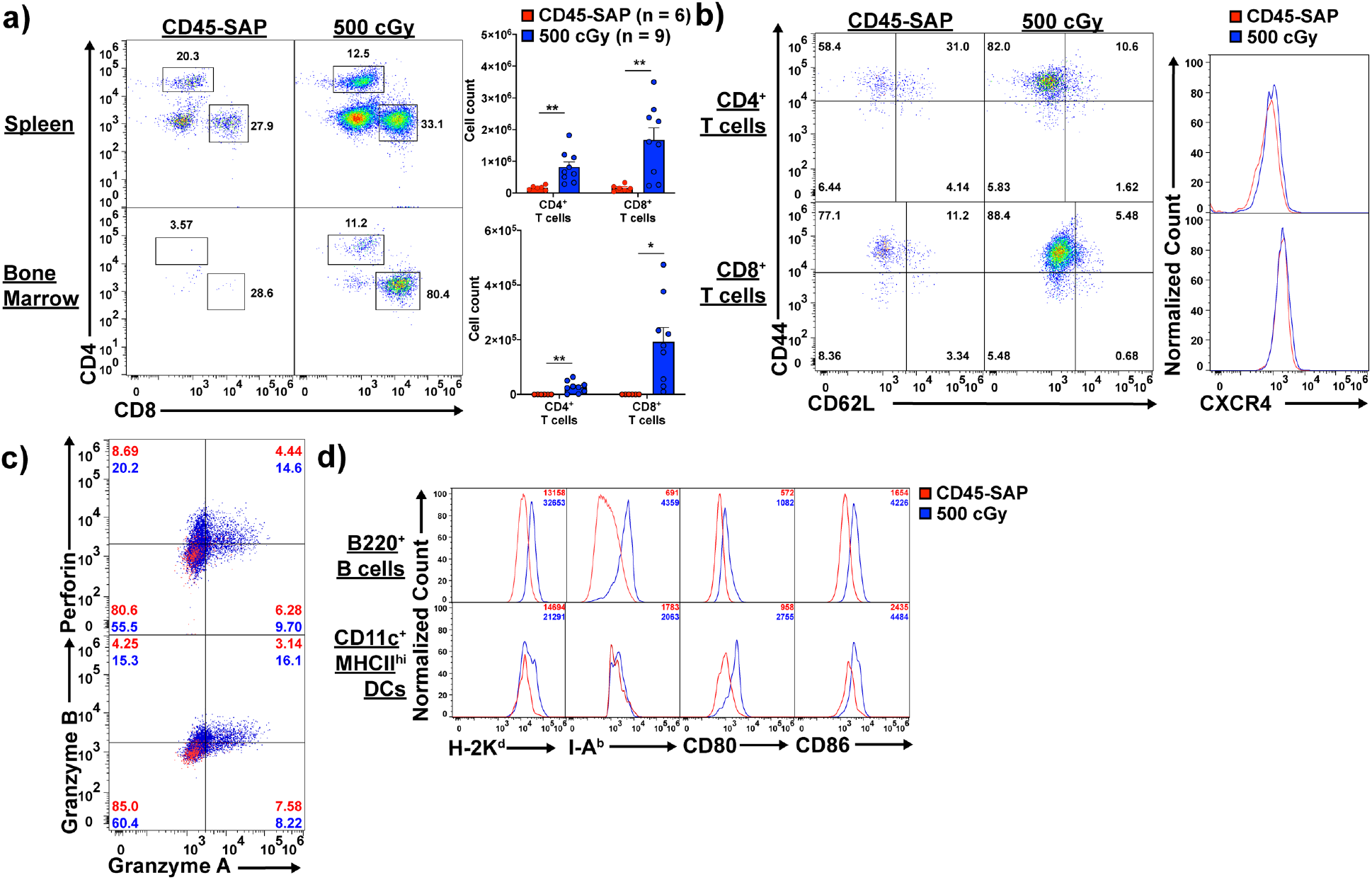
Irradiation, but not CD45-SAP, promotes alloreactive T cell expansion, effector function, and bone marrow infiltration. **(a)** Absolute counts of donor-derived (H-2K^b+^/^d-^) CD4^+^ and CD8^+^ T cells in spleens and bone marrows of CB6F1 mice conditioned with 500 cGy total body irradiation (TBI) or CD45-SAP at 7 days post-infusion of allogeneic B6 splenocytes. (b) Cell surface phenotyping of donor T cells harvested from spleens of TBI-versus ADC-conditioned mice. (c) Intracellular staining of donor T cells harvested from spleens of TBI-versus ADC-conditioned mice for CD8^+^ T cell cytolytic granule enzymes. (d) Cell surface phenotyping of the recipient (H-2K^b+/d+^) APC compartment in spleens of TBI-versus ADC-conditioned mice. For panels (b) and (c), inset numbers indicate the percent of events in each quadrant; for (d), inset numbers are MFIs. FACS plots are from one representative mouse obtained across 2 (CD45-SAP) or 3 (500 cGy) experiments; data points and error bars represent mean ± SEM. For statistical comparisons: ns = not significant, * = p<0.05, ** = p<0.01, *** = p<0.001, **** = p<0.0001.

## DISCUSSION

Toxicities from chemotherapy- and radiation-based conditioning remain a major obstacle to the broader application of HSCT for the treatment of hematopoietic diseases, particularly for elderly or infirmed patients. Reduced-intensity conditioning provides one way to extend HSCT to patients unable to tolerate more severe conditioning,^47^ and is a reasonable approach to treat non-malignant diseases or for autologous gene therapy, for which mixed donor chimerism may be sufficient for cure. However, for AML, reduced-intensity conditioning more poorly ablates residual malignant cells, potentially leading to relapse. In this setting, relapse control becomes more reliant on GvL effects^48, 49^, which is inextricably linked to GvHD development. These issues illustrate the complexity of managing treatment-related toxicity and relapse outcomes to achieve optimal outcomes for patients receiving allo-HSCT for leukemia.

Antibody-based HSCT conditioning presents a way to more favorably balance toxicity and therapeutic efficacy. By simultaneously targeting the stem cell compartment and malignant cells, the therapeutic goals of HSCT can hypothetically be achieved with toxicities largely confined to the hematopoietic system. Indeed, recent work in murine^17, 18, 22, 23, 25, 50^, non-human primates^51, 52, 53^, and early human trials^54^ have demonstrated feasibility and limited toxicities of antibody and ADC-based therapies alongside high efficacy in depleting recipient HSCs and/or malignant cells. Optimization of these strategies for allo-HSCT and translation to human clinical trials will benefit from a greater mechanistic understanding of how they modulate donor and recipient immunity, with significant implications for treating high-risk malignancies like AML.

In the present study, we used mouse allo-HSCT models to identify ADC-based conditioning regimens able to achieve robust donor engraftment, and to understand the immunobiology underlying their effect. T cells were the primary immune barriers to miHA- and MHC-mismatched HSCT, as engraftment was achievable by combining CD45-SAP with pan-TCD. The transient donor chimerism we often encountered in the MHC-mismatched model could be explained by incomplete T cell elimination at all tissue sites, even with prolonged depleting antibody treatment. This issue is particularly relevant for developing thymocytes, which not only require higher antibody doses than peripheral T cells for depletion^55^, but also include a CD4^-^CD8^-^ subset unable to bind αCD4 or αCD8 antibodies. Upon maturation and thymic egress, these cells could mediate alloreactivity in the periphery. Given the high frequency of T cells estimated to be alloreactive^56^, even a small residual population of functional host T cells could reject donor HSCs.

The use of baricitinib in combination with CD45-SAP as conditioning improved significantly upon the shortcomings of pan-TCD, suppressing host T and NK cell responses to enable robust, high-level, multilineage engraftment without requiring prolonged antibody depletion. As baricitinib relatively spares JAK3^57^, its inhibitory activity against T and NK cells may result from antagonism of JAK1, which associates with the common beta chain (CD122) used by the IL-2 and IL-15 receptors. Interference with IL-15 signaling led to the poorer *in vitro* proliferation, survival and function we observed with baricitinib-treated NK cells, and an analogous mechanism impacting IL-2 signaling may affect baricitinib-treated T cells. Future studies utilizing inhibitors targeting individual JAKs will help further dissect the mechanisms of baricitinib effect on T and NK cell biology and identify the relevant signaling pathways.

While daily baricitinib monotherapy was sufficient to permit engraftment in miHA-mismatched HSCT, co-administration of pre-HSCT TCD or continuous infusion of baricitinib was required for the MHC-mismatched model, consistent with our expectation that more durable immunosuppression is necessary with increasing degrees of MHC disparity. Indeed, in our preliminary experiments with fully MHC-mismatched allo-HSCT (BALB/c→B6), continuous baricitinib plus CD45-SAP conditioning gave ~25% engraftment success (not shown), considerably lower than the ~70% engraftment rate for the MHC-mismatched model described herein. Continued optimization of our conditioning regimens aims to achieve robust, high-level engraftment in fully haploidentical (F1-to-F1) and fully MHC-mismatched allo-HSCT. ADCs with alternative toxic payloads able to produce greater myeloablation and lymphodepletion, or immunosuppressive treatments able to synergize with baricitinib, are potential avenues to improvement in this regard.

Our finding that daily baricitinib synergized with pre-HSCT TCD in our MHC-mismatched model was in some respects surprising. That baricitinib could substitute for post-HSCT TCD is consistent with our hypothesis that T cell inhibition is crucial to its activity. However, this strategy succeeded in mice whose CD4^+^ and CD8^+^ T cells were ablated by pre-HSCT T cell depletion. While this could simply reflect baricitinib inhibiting the few cells surviving TCD, other explanations are possible, such as modulation of APC function or alterations of thymic selection or egress. Aside from NK cell depletion, we did not observe gross immunologic alterations in mice acutely treated with baricitinib, yet there may be more subtle effects that impact allograft tolerance or effects which become apparent only with chronic treatment. Deeper immunophenotyping and transcriptomic profiling in baricitinib-treated mice, focusing on differences between daily versus continuous baricitinib administered either acutely or chronically, is central to rationally designing and optimizing allo-HSCT conditioning with baricitinib.

Finally, infusion of allogeneic T cells did not elicit pathogenic alloreactivity in CD45-SAP conditioned mice due to poor donor cell expansion, effector function, and target organ infiltration. This outcome likely reflects poorer priming of the donor alloresponse by innate immune signals in ADC-conditioned mice, signals that are more abundantly generated by TBI-induced injury^58, 59^. Given the large numbers of T cells in peripheral blood-mobilized stem cell preparations, an ADC-based conditioning regimen that minimizes collateral tissue damage might prevent amplification of T cell alloreactivity leading to GvHD. However, donor T cells unable to elicit GvHD may also be unable to mount a GvL response, which could offset the benefit of reduced GvHD with a greater risk of leukemia relapse. Murine leukemia models utilizing allo-HSCT with ADC-based conditioning would provide a relevant preclinical platform on which to integrate *in vivo* studies of engraftment, GvHD and GvL effects and understand the underlying biology.

In conclusion, the studies presented herein exemplify the promise of immunotherapy to provide safe, effective conditioning for HSCT. Importantly, our studies provide insights to the unique immunobiology of ADC-conditioned allo-HSCT and an experimental foundation on which further basic and translational investigations can be conducted.

## METHODS

### Mice

Mice were handled in accordance with an animal protocol approved by the Institutional Animal Care and Use Committee (IACUC) at Washington University School of Medicine. The following strains were used in our studies: C57BL6/J, BALB/cJ, DBA/2J, CB6F1/J (C57BL6/J x BALB/cJ F1), C57BL/6-Tg(UBC-GFP)30Scha/J (B6-GFP), CByJ.SJL(B6)-*Ptprc^a^*/J (BALB-Ly5.1). IFNγR^-/-^ mice were provided by Dr. Herbert Virgin (Department of Pathology and Immunology, Washington University). All mice were bred within specific-pathogen free colonies at Washington University School of Medicine or purchased from Jackson Laboratories (Bar Harbor, ME) and maintained on *ad libitum* water and standard chow (LabDiet 5053; Lab Supply, Fort Worth, TX). Age and gender-matched mice were used for all experiments, with all animals aged 6-12 weeks old; no selection was applied to assign mice to experimental treatment groups. For all experiments involving lethal irradiation, mice received trimethoprim-sulfamethoxazole (SulfaTrim, 5 mL per 250 mL drinking water) for two weeks beginning two days prior to irradiation. For retroorbital injections, mice were anesthetized with 3% isoflurane in O2 delivered by vaporizer at a flow rate of 1 L/min. For survival surgery procedures (skin grafting and osmotic pump implantation), mice were anesthetized via intraperitoneal injection of 80-100 mg/kg ketamine plus 5-10 mg/kg xylazine. Prior to first incision, the surgical site was shaved, disinfected, and draped in sterile fashion. Skin closure was done using 9 mm autoclips and buprenorphine (0.1 mg/kg) was provided for post-operative analgesia.

### Mouse tissue preparation

Spleens, lymph nodes, or thymi harvested from euthanized mice were processed into single-cell suspensions by gentle homogenization with a syringe plunger through a 70 μm filter in PBS containing 0.5% BSA and 2 mM EDTA (Running buffer). Bone marrow was harvested from femurs and tibias by centrifugation as previously described^60^. Mouse peripheral blood samples were drawn from the facial vein using Goldenrod 5 mm animal lancets (Medipoint; Mineola, NY) and collected into K3EDTA-coated tubes (BD). Erythrocytes were removed from all mouse tissue specimens using ammonium chloride-potassium bicarbonate (ACK) lysis.

### Cell culture and *in vitro* assays

Primary murine T cells were grown in R10 media - RPMI plus 10% fetal bovine serum (FBS; R&D Systems, Minneapolis, MN) supplemented with GlutaMAX (Gibco) and penicillin/streptomycin (Gibco) - at 37°C/5% CO_2_. For mouse NK cell cultures, R10 media was supplemented with 10 mM HEPES, 0.1 mM non-essential amino acids, 1 mM sodium pyruvate and 55 μM 2-mercaptoethanol (K10 medium). YAC-1 cells for NK cell cytotoxicity assays were obtained from ATCC, tested negative for *Mycoplasma*, and maintained in R10 media. Human peripheral blood mononuclear cells (PBMCs) were harvested from leukoreduction chambers by Ficoll density gradient centrifugation and cryopreserved.

For *ex vivo* stimulation of primary mouse T cells, 1 x 10^5^ T cells purified from spleen and lymph nodes with the EasySep Mouse T cell Isolation Kit (Stem Cell Technologies; Vancouver, BC, Canada) were cultured with 4 x 10^5^ T-depleted splenocytes in 96-well round bottom plates in R10 media with 1 μg/mL αCD3. Supernatants were collected for cytokine analysis after 24 hours incubation, and cultured cells were analyzed for expansion by flow cytometry at 72 hours.

For primary NK cell assays, splenic NK cells were enriched to >80% purity using the EasySep Mouse NK Cell Isolation Kit (Stem Cell Technologies), and 1.0-2.5 x 10^4^ NK cells were cultured in K10 media along with either IL-15 alone (1-100 ng/mL; BioLegend), or a cocktail of IL-12 (10 ng/mL; BioLegend), IL-15 (10 ng/mL), and IL-18 (50 ng/mL; BioLegend)^36^. For cytotoxicity assays, purified NK cells were stimulated with 100 ng/mL IL-15 for 48 hours, washed twice to remove cytokines, then incubated for 4 hours at multiple effector-to-target ratios with a fixed number of YAC-1 target cells (2000 YAC-1 cells per well of a 96 well, round-bottom plate) with or without baricitinib. YAC-1 cell death was assessed with Zombie viability dye staining and analyzed by flow cytometry.

For colony forming unit (CFU) assays, murine whole bone marrow was resuspended at 10X final concentration in IMDM + 2% FBS with single doses or concentration series of ADC or control conjugate. Then, 300 μL of each suspension was diluted into 3 mL complete methylcellulose medium (R&D), vigorously vortexed, and 1.1 mL of the resulting methylcelluose suspension was plated in duplicate and incubated 12 days at 37°C.

### Complete blood counts (CBC)

CBC analysis was performed using a Hemavet 950 analyzer (Drew Scientific). Total white blood cell (WBC) count plus differential, hematocrits (Hct), and platelet (PLT) counts were obtained. Reference ranges were as follows: WBC 1.8-10.7 x 10^3^ cells per μL, Hct 35.1-45.4%, PLT 592-2972 x 10^3^ cells/μL. Absolute counts of circulating leukocyte subsets were calculated by multiplying the WBC by the frequency of each specific cell type as measured by flow cytometry.

### Flow cytometry

Flow cytometry was performed with a Beckman Coulter Gallios instrument equipped with Kaluza acquisition software. *Post hoc* compensation and data analysis were done using FlowJo version 10.7 (Treestar; Ashland, OR). All flow cytometry reagents are listed in Supplementary Table 1. For routine preparation of fresh, unfixed samples, single cell suspensions were stained with fluor-conjugated antibodies to surface antigens in 100 μL Running buffer at room temperature for 15-20 minutes. For staining with biotinylated antibodies, samples were first incubated with biotinylated antibody, washed, then stained with fluor-conjugated streptavidin plus any other fluor-conjugated antibodies as above. For viability staining of fresh samples, 7-aminoactinomycin D (7-AAD; BioLegend, San Diego, CA) was added at 1 μg/mL immediately before analysis. For intracellular cytokine and cytotoxic granule staining, cells were stained with Zombie fixable viability dye (BioLegend; 1:400 final dilution) in PBS for 15 minutes, then stained 15 minutes for surface markers and fixed for 20 minutes with 4% paraformaldehyde (PFA) in PBS (BioLegend). Cells were then permeabilized with 0.5 % saponin in Running buffer and stained for intracellular markers. FoxP3 staining was done using the FoxP3/Transcription Factor Staining Buffer Set per the manufacturer’s instructions (eBioscience).

### Phosphoflow analysis

For phospho-Stat1 analysis, whole blood from baricitinib- or vehicle-treated mice was stimulated for 15 minutes with 100 ng/mL murine IFNg at 37°C, then immediately fixed with 1 mL Lyse/Fix Buffer (BD) for 10 minutes at 37°C. For phospho-Stat3 analysis, cryopreserved human PBMC were thawed and rested overnight at 37°C in R10, stimulated with 100 ng/mL human IL-6 for 15 minutes at 37°C in the presence of baricitinib or vehicle (0.1% DMSO), then fixed in 4% PFA in PBS. For phospho-Stat5 analysis, purified B6 mouse splenic NK cells were incubated for 30 minutes with 100 ng/mL IL-15 in K10 medium in the presence of baricitinib or vehicle, then fixed in 4% PFA.

After stimulation and fixation, all samples were permeabilized in ice-cold Perm Buffer III (BD) and held at −20°C overnight. Samples were then washed thrice with Running buffer and stained for phospho-Stat molecules. For phospho-Stat1, samples were incubated 1 hour at room temperature with primary rabbit anti-phospho Stat1 (Y701, Cell Signaling Technology #9167, clone 58D6), then washed and stained 1 hour with Alexa Fluor 647-conjugated anti-rabbit secondary antibody (Cell Signaling Technology #4414). For phospho-Stat3, samples were stained with anti-human CD4 and anti-phospho Stat3 (Y705, BD Biosciences) for 1 hour at room temperature. For phospho-Stat5, samples were stained with anti-NK1.1 (BioLegend) and anti-phospho Stat5 (Y694, BD Biosciences).

### Cytokine analysis

Cytokine concentrations in culture supernatant or mouse plasma were measured with the LegendPLEX Inflammation Panel (13-plex) or the Mouse Th1 Panel (5-plex) per the manufacturer protocols (BioLegend). Quantification was done using LegendPLEX software v8.0 for Windows. If a cytokine concentration was too low to be quantified, the sample was assigned the value of the lower limit of quantitation. For intracellular IFNγ analysis of splenic NK cells, cells were cultured in K10 media with or without cytokine stimulation for 15 hours, with 5 μg/mL Brefeldin A (BioLegend) added to each well for the last 2.5 hours. After this incubation period, cells were fixed, saponin-permeabilized, and stained as described above.

### Preparation of saporin antibody-drug conjugates (ADC)

Saporin conjugated to streptavidin (sAV-SAP; Advanced Targeting Systems, San Diego, CA) was used to indirectly couple biotinylated antibodies to saporin to generate the ADCs used in this study. The average saporin-to-streptavidin ratio was 2.4, yielding an effective molecular weight of 127 kDa. A total molecular weight of 287 kDa (127 kDa for sAV-SAP + 160 kDa for IgG) was used for conversions between molar and mass concentrations.

Saporin-linked ADCs targeting murine CD45.2 (CD45-SAP) and cKit (cKit-SAP) were generated by incubating biotinylated anti-mouse CD45.2 (clone 104, BioLegend) or biotinylated anti-mouse cKit (clone 2B8, BioLegend) with sAV-SAP in a 1:1 molar ratio for 15 minutes at room temperature. Afterwards, ADCs were diluted to their final concentration in endotoxin-free PBS (Sigma-Millipore) and injected intravenously via the retroorbital sinus (100-150 μL per injection). Prior to ADC generation, sodium azide and endotoxin were removed from the biotinylated antibodies with Zeba desalting spin columns and High-Capacity Endotoxin Removal spin columns (ThermoFisher) per the manufacturer’s instructions, then filter-sterilized using an 0.22 μm PES syringe filter.

For control experiments in which free antibody and free sAV-SAP were administered together, non-interaction of these two components was ensured by using non-biotinylated antibodies and sAV-SAP whose biotin-binding sites were occupied by an irrelevant biotinylated 11-mer peptide (BLANK Streptavidin-SAP, Advanced Targeting Systems). For experiments in which free antibody or sAV-SAP were administered alone, the equivalent mass of each component alone in the ADC was administered to each mouse (i.e., the doses of CD45.2 antibody and sAV-SAP corresponding to a CD45-SAP dose of 75 μg would be 41.8 μg and 33.2 μg, respectively). To avoid interference by cKit-SAP, bone marrows analyzed by flow cytometry were stained for c-Kit using clone ACK2, which does not compete for binding with clone 2B8.

### Hematopoietic stem cell transplantation with ADC conditioning

Mice were injected with CD45-SAP or cKit-SAP at doses indicated in each figure at 7 days pre-transplant (d-7). In general, 3 mg/kg (75 μg) CD45-SAP, and 0.4 mg/kg or 2 mg/kg (10 or 50 μg, respectively) cKit-SAP were used. On transplant day (d0), mice received 10 x 10^6^ whole donor bone marrow cells via the retroorbital injection. Mice conditioned with saporin-ADCs did not receive antibiotic prophylaxis.

For serial transplantation experiments, mice received a single dose of lethal irradiation (1100 cGy for B6 mice, 950 cGy for DBA/2 mice) from a Mark I Model 30 irradiator (J.L. Shepherd and Associates, ^137^Cs source, 73.69 cGy/min as tested on 1/1/2020) and transplanted with 10 x 10^6^ whole bone marrow cells from primary transplant recipients 8-16 hours post-irradiation.

### *In vivo* lymphocyte depletion

Antibodies for *in vivo* T and NK cell depletion and isotype controls were obtained from BioXcell (West Lebanon, NH) in azide-free, low-endotoxin formulations confirmed to be murine pathogen-negative for (InVivoPlus grade). CD4^+^ and CD8^+^ T cell depletion was done using clones GK1.5 and YTS169.4, respectively, and NK cell depletion done using clone PK136. Mouse IgG2ak (clone C1.18.4) and rat IgG2bk (clone LTF-2) were used as isotype controls. All antibodies were administered intraperitoneally at 250 μg per dose following the schema for each experiment. Depletion of target cell populations was routinely confirmed by flow cytometry of peripheral blood immediately prior to HSCT; to avoid interference with depleting antibodies, CD4^+^ T cells were stained for flow cytometry with clone RM4-4, CD8^+^ T cells with clone 53-6.7, and NK cells with a combination of CD3, CD49b (DX5) and NKp46.

### Daily infusion with JAK inhibitors

The selective Janus kinase 1 and 2 (JAK1/2) inhibitors baricitinib (LY3009104, INCB028050) and ruxolitinib (INCB18424) were obtained from MedChemExpress (Monmouth Junction, NJ). For subcutaneous administration, baricitinib was dissolved in 100% DMSO at 20 mg/mL and stored at −20°C. Immediately prior to injection, these DMSO stocks were thawed, diluted 1:10 in PBS, and injected at 200 μl/mouse subcutaneously (400 μg daily dose). For HSCT experiments, mice were treated with baricitinib or vehicle (10% DMSO in PBS) for a total of 25 days, beginning at d-3 relative to transplant and ending at d+21.

### Osmotic pump administration of JAK inhibitors

ALZET subcutaneous osmotic pumps (Model 2004) were used to continuously deliver JAK1/2 inhibitors to mice for 28 days at a rate of 0.25 μL/hour (6 μL/day). A vehicle of 50% dimethyl sulfoxide (DMSO)/50% polyethylene glycol 400 (PEG-400) was used for all experiments. JAK1/2 inhibitors were prepared at 2X concentration (133.3 mg/mL) in 100% DMSO then diluted to 1X with an equal volume of PEG-400 (66.7 mg/mL, 400 μg total daily dose). Prepared compounds were then loaded into osmotic pumps per the manufacturer’s instructions and surgically implanted in accordance with our IACUC-approved protocol.

### Skin grafting

Surgical engraftment of donor ear skin to recipient mice was performed as described^61^. Briefly, donor BALB/cJ and DBA/2J mice were euthanized, and ear skin was harvested and held in ice-cold PBS in preparation for transplant. Skin graft recipients were DBA/2J mice that either successfully engrafted with BALB-Ly5.1 bone marrow (BALB-DBA mixed chimeras) or those that failed to engraft. /mice mice were prepped for survival surgery as described above and had a small patch of dorsal skin resected and replaced with donor ear skin. Recipients were then bandaged, single-housed, and monitored for 4 days to ensure the bandage and graft bed remained undisturbed. Bandages were then removed, and graft recipients monitored daily for signs of rejection (scabbing, wound contraction).

### Graft-versus-host alloreactivity model

A parent-to-F1 adoptive transfer model was used to study T cell alloreactivity *in vivo*, as previously described^46^. In this model, irradiated CB6F1 mice receiving allogeneic B6 splenocytes develop immune-mediated bone marrow aplasia, with lethality occurring approximately 3 weeks post-T cell infusion. Recipients were conditioned either with CD45-SAP 7 days before adoptive transfer, or with 500 cGy irradiation delivered 8-16 hours pre-adoptive transfer. After conditioning, recipients were infused with 25 x 10^6^ splenocytes from B6 mice. As negative controls, irradiated CB6F1 mice were treated with CB6F1 (syngeneic) splenocytes, and non-conditioned CB6F1 mice were treated with B6 splenocytes. Clinical scoring of mice was done based on a 10-point scale (0-2 points each for posture, activity, fur ruffling, weight loss, and skin lesions), with higher scores indicating worse disease as previously described^62^. No mice in these studies received antibiotic prophylaxis.

### *In vivo* mixed leukocyte reactions (MLR)

Recipient mice were infused with 2-3 x 10^6^ purified donor T cells that were labeled with 5 μM CFSE (BioLegend) as previously described^63^. Recipients were euthanized at 72 h post-T cell infusion and splenocytes analyzed for CFSE dilution of the infused donor T cells.

### Histopathology

Femurs (for bone marrow histology) were preserved in neutral buffered formalin (PBS plus 3.7% formaldehyde) and incubated at room temperature with gentle rocking for at least 48 hours. Fixed samples were submitted to the Washington University Department of Comparative Medicine Animal Diagnostic Lab for decalcification and preparation of formalin-fixed paraffin embedded sections and staining with hematoxylin and eosin. A trained veterinary pathologist who was blinded to the experimental treatments provided descriptive reports of any pathological findings.

### Data analysis and statistics

Sample size determinations and analysis parameters were based on general guidelines for laboratory animal research^64^. Data for all experiments were compiled and statistically analyzed using GraphPad Prism version 8.0 for Mac. IC50 values for cytotoxicity studies were calculated by curve-fitting the dose response data to a three- or four-variable inhibition model. The Shapiro-Wilk test for normality was used to assess conformity of each dataset to a normal distribution. For comparison of two normally distributed datasets, unpaired, two-tailed Student’s *t* tests with Welch’s correction (no assumption of equal variance between groups) were used; if either dataset was not normally distributed, the Mann-Whitney *U* test was used instead. Survival analysis was done with the Mantel-Cox log-rank test. For comparisons of CBC values with the lower reference limit, a one-sample *t* test was used. The criterion for statistical significance for all comparisons was p ≤ 0.05.

## Supporting information

Supplementary Figures 1-9 plus legends

Supplementary Table 1 - Flow reagent list

## ACKNOWLDGMENTS

This study was funded by NIH/NCI R35CA210084 (J.F.D), NIH/NCI Leukemia SPORE grant (P50CA171963, J.F.D.) and Leukemia SPORE Career Enhancement Award (P50CA171063, S.P.P.), an American Society of Transplantation and Cellular Therapy New Investigator Award (S.P.P.), an NCI Research Specialist Award (R50CA211466, M.P.R), a donation from Bigelow Aerospace (J.F.D.), and awards from Gabrielle’s Angel Foundation for Cancer Research (S.P.P.). J.C was supported by The Amy Strelzer Manasevit Research Program (Be The Match Foundation and National Marrow Donor Program) and Alvin J. Siteman Cancer Center Siteman Investment Program (supported by the Foundation for Barnes-Jewish Hospital Cancer Frontier Fund, National Cancer Institute Cancer Center Support Grant, P30CA091842, and Barnard Trust). B6 mice for *in vitro* T and NK cell assays were a kind gift from P.M. Allen. We thank J. Eissenberg for helpful discussions and feedback, and P.M. Allen for critical review of the manuscript.

## AUTHOR CONTRIBUTIONS

S.P.P, P.G.R., M.L.C., M.P.R., and J.F.D conceived and designed the research, S.P.P. and J.K.R. conducted the experiments, J.C. contributed data, S.P.P. performed data analysis, and S.P.P and J.F.D. wrote the manuscript. All authors provided scientific and technical feedback on the work and approved the final manuscript for submission.

## COMPETING INTERESTS

The authors declare the following competing interests:

**S.P.P.:** None to declare

**J.K.R:** None to declare

**J.C.:** Consultancy – Daewoong Pharmaceutical; Research support: Mallinckrodt Pharmaceuticals, Incyte Corporation.

**P.G.R.:** None to declare

**M.L.C.:** None to declare

**M.P.R.:** None to declare

**J.F.D.:** Consulting/Advisory Committees – Rivervest; Research support – Macrogenics, BioLine, NeoImmuneTech, Incyte Corporation; Ownership Investment – Magenta Therapeutics, Wugen.

## ABBREVIATIONS

7-AAD: 7-aminoactinomycin D
Allo-HSCT: allogeneic hematopoietic stem cell transplantation
ACK: Ammonium chloride-potassium bicarbonate
ADC: antibody-drug conjugate
AML: acute myeloid leukemia
APC: antigen presenting cell
BSA: bovine serum albumin
CBC: Complete blood count
CD: Cluster of differentiation
CFU: colony forming unit
FBS: fetal bovine serum
EDTA: ethylenediaminetetraacetic acid
FACS: fluorescence-activated cell sorting
GFP: green fluorescent protein
vGHD: graft-versus-host disease
GvL: graft-versus-leukemia
Hct: hematocrit
HEPES: *A-2*-hydroxyethylpiperazine-*N′-2*-ethanesulfonic acid
HSC/HSCT: hematopoietic stem cell/hematopoietic stem cell transplantation
IFN: Interferon
Ig: immunoglobulin
IL: Interleukin
IMDM: Iscove’s Modified Dulbecco’s Media
JAK: Janus kinase
LK: Lineage^-^Sca1^-^cKit^+^
LSK: Lineage^-^Sca1^+^cKit^+^
miHA: minor histocompatibility antigen
MHC: major histocompatibility complex
MLR: mixed leukocyte reaction
PLTS: platelets
PBS: phosphate buffered saline
Running buffer: PBS + 0.5% BSA + 2 mM EDTA
NK: natural killer cell
RAG: recombinase activating gene
RPMI-1640: Roswell Park Memorial Institute-1640 media
sAV: streptavidin
SAP: saporin-conjugated
Stat: signal transducer and transactivator
TBI: Total body irradiation
TCD: T cell depletion/depleted
TNF: Tumor necrosis factor
WBC: white blood cells

