## Supplementary Figures 1-9 plus legends for "Antibody-drug conjugates targeting CD45 plus Janus kinase inhibitors effectively condition for allogeneic hematopoietic stem cell transplantation"

### SUPPLEMENTARY FIGURES AND LEGENDS

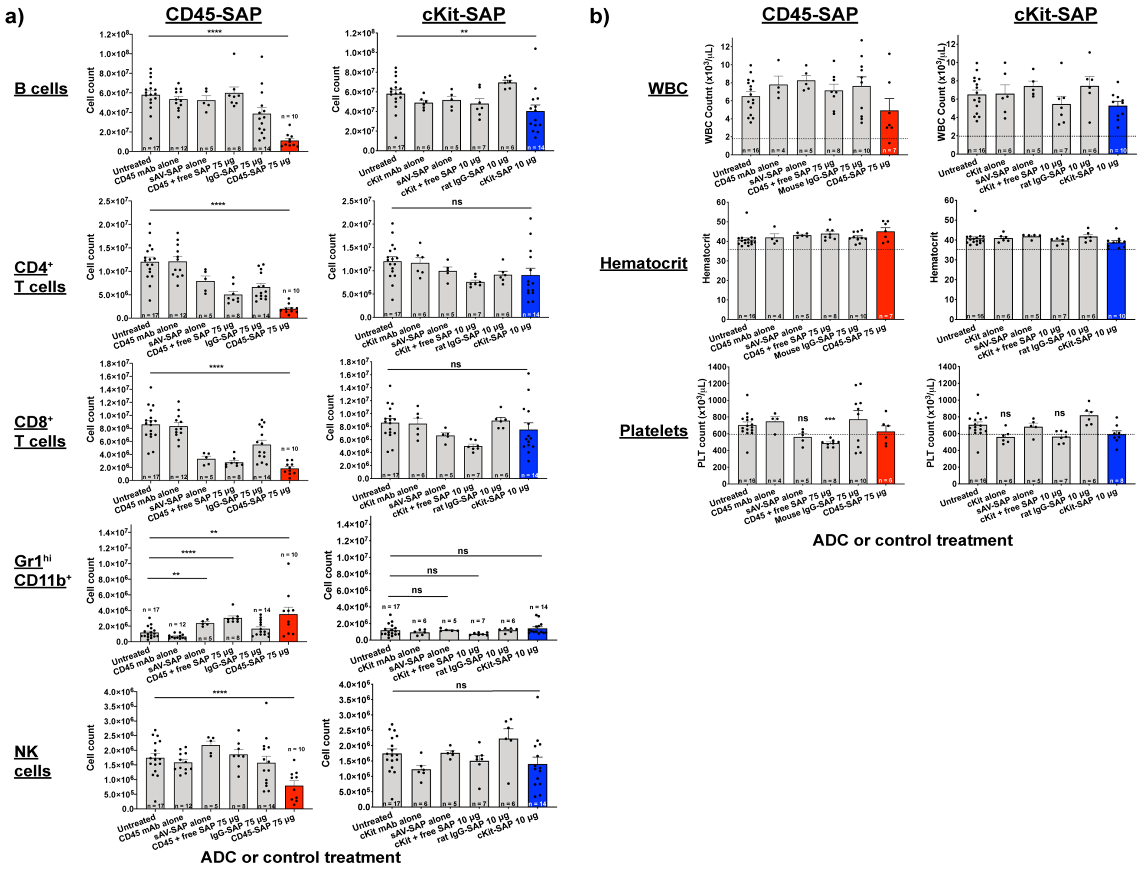


**Supplementary Figure 1. Acute hematologic effects of CD45-SAP and cKit-SAP conditioning. (a and b)** Absolute leukocyte counts by subset in mouse spleen (a) and CBCs (b) 7 days after administration of CD45-SAP, cKit-SAP, or control ADCs. Dotted lines for the CBC assays indicate the lower reference limits; groups whose means were statistically below the lower reference limit are indicated. Please note that the same cohort of untreated mice was used to compare with mice in the CD45-SAP and cKit-SAP groups. Data points and error bars represent mean ± SEM. For statistical comparisons: ns = not significant, * = $p < 0.05$, ** = $p < 0.01$, *** = $p < 0.001$, and **** = $p < 0.0001$.

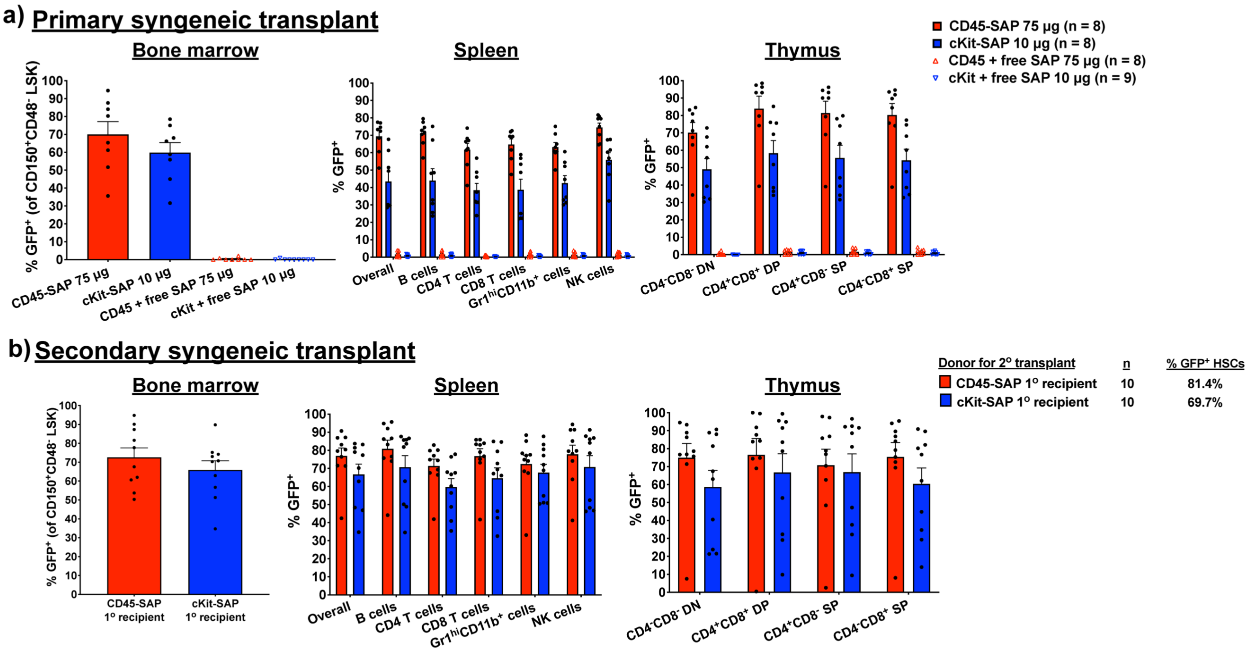


**Supplementary Figure 2. Donor engraftment in lymphoid organs of primary and secondary syngeneic HSCT**. **(a)** Donor chimerism in bone marrow, spleen and thymus 6 months after primary syngeneic (B6-GFP→B6) HSCT; these data correspond to the peripheral blood donor chimerism data shown in Figure 1c. For thymus, DN = double negative. DP = double positive, SP = single positive. **(b)** Donor chimerism in bone marrow, spleen, and thymus 4 months after secondary transplantation of bone marrow obtained from B6-GFP→B6 primary recipients to a new cohort of lethally-irradiated B6 mice; these data correspond to the peripheral blood donor chimerism data shown in Figure 1e. The percentages of primary recipient-derived, $GFP^+$ HSC that were infused into the secondary recipients is provided. Data points and error bars represent mean ± SEM.

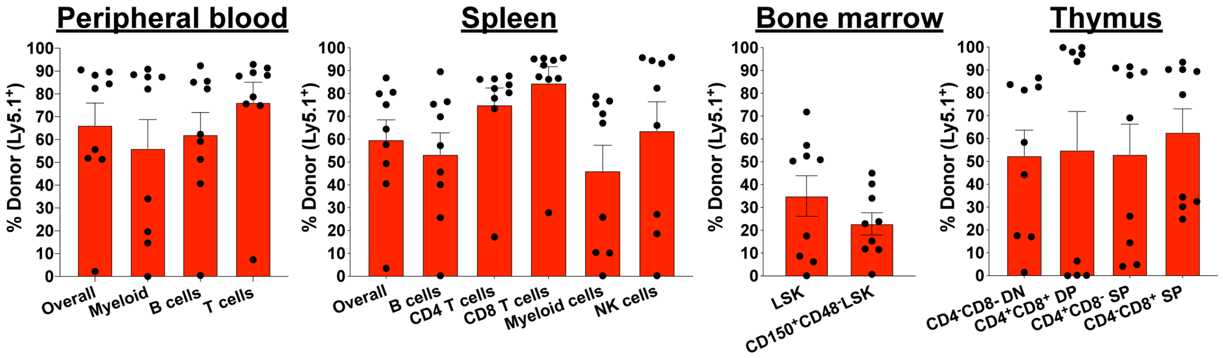


**Supplementary Figure 3. Serial transplantation in CD45-SAP conditioned, miHA-mismatched alloHSCT.** $10^7$ whole bone marrow cells isolated from BALB/c-Ly5.1→DBA/2 mixed chimeras were infused to a new cohort of lethally-irradiated DBA/2 mice. The percentage of primary recipient-derived, $Ly5.1^+$ HSCs that were transferred to secondary recipients was ~76%. Donor chimerism in peripheral blood, bone marrow, spleen, and thymus of secondary recipients were assessed at 4 months post-transplant. Data points and error bars represent mean ± SEM of 9 mice from one experiment.

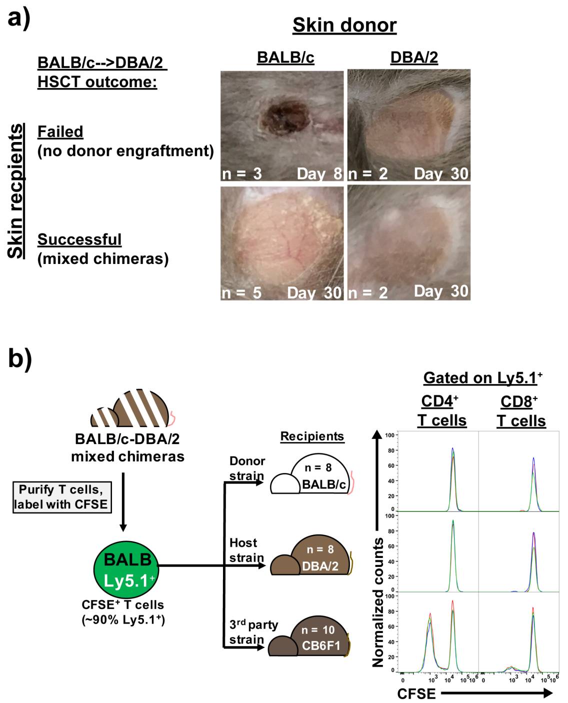


**Supplementary Figure 4. BALB/c→DBA/2 mixed chimeras are cross-tolerant to donor and recipient-derived antigen. (a)** BALB/c or DBA/2 ear skin was surgically engrafted at 6 months post-HSCT to CD45-SAP-conditioned, BALB/c→DBA/2 recipients that had either rejected or successfully engrafted donor HSC. Skin grafts were monitored for 30 days for signs of rejection; inset text indicates the number of skin graft recipients analyzed across two experiments (lower left) and the time post-skin graft when images were acquired (lower right). **(b)** *In vivo* MLRs in which CFSE-labeled T cells from BALB/c-DBA/2 mixed chimeras were infused to new cohorts of BALB/c, DBA/2, or CB6F1 mice. The indicated numbers of recipient mice per group were analyzed across two experiments, with CFSE histograms from three representative mice per group presented.

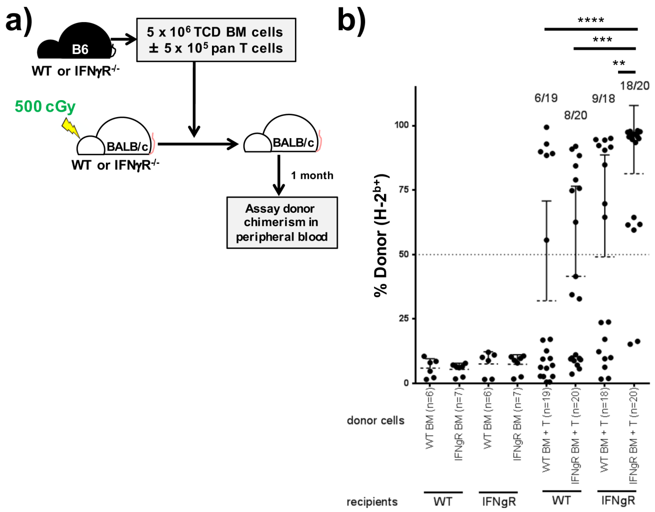


**Supplementary Figure 5. Deficiency of IFNγ signaling permits engraftment of fully-mismatched HSC in a model of reduced-intensity alloHSCT. (a)** Schema for reduced-intensity conditioning HSCT model in which WT or IFNγR$^{-/-}$ BALB/c recipients were sublethally irradiated then transplanted with bone marrow with or without T cells from WT or IFNγR$^{-/-}$ B6 mice. **(b)** Peripheral blood chimerism at 1 month post-HSCT. Results are pooled from three independent experiments; the frequency of mice in each group with greater than 50% donor chimerism is indicated above each dataset. For statistical comparisons, * = $p < 0.05$, ** = $p < 0.01$, *** = $p < 0.001$, and **** = $p < 0.0001$.

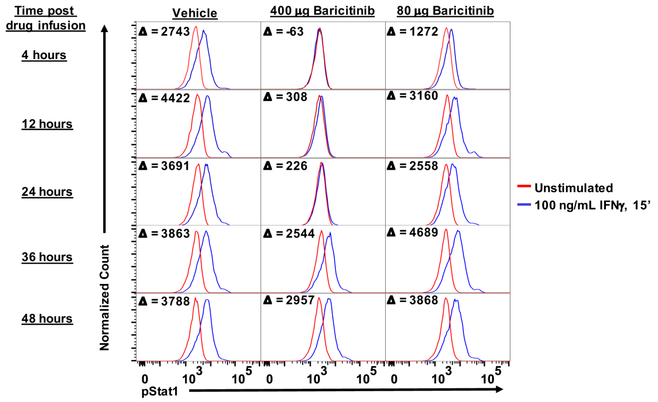


**Supplementary Figure 6. Pharmacodynamics of subcutaneously-administered baricitinib.** B6 mice received a single subcutaneous injection of baricitinib (400 µg or 80 µg) or vehicle, and Stat1 phosphorylation of whole blood leukocytes ($CD45^+$ gated) in response to IFNγ stimulation (100 ng/mL, 15 minutes) was assayed at the indicated times post-drug infusion. Data from a single mouse in each treatment group are displayed and are representative of 2 (vehicle group) or 4 (baricitinib groups) mice analyzed over two experiments. Inset numbers are the difference in MFI between IFNγ-stimulated and unstimulated samples.

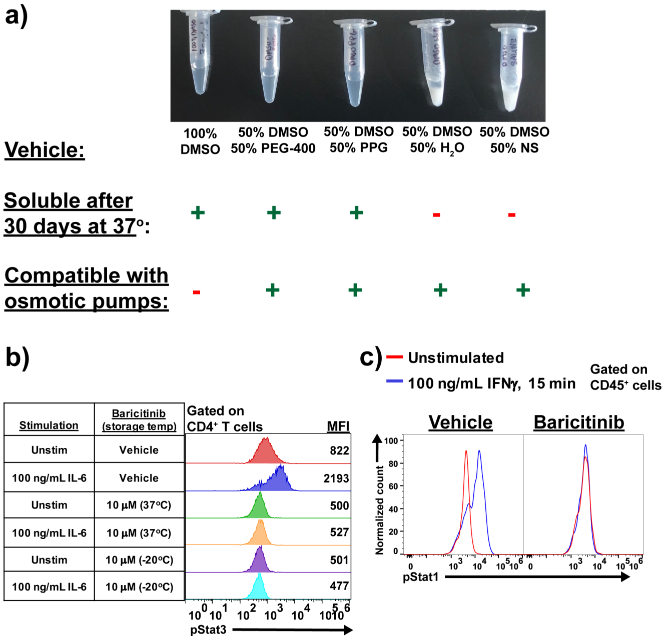


**Supplementary Figure 7. Baricitinib is compatible with *in vivo* drug delivery via osmotic pump. (a)** Solubility testing of baricitinib (70 mg/mL) in various test solvents after 30 days incubation at 37°C. Vehicle compatibility is as per the manufacturer. PEG = polyethylene glycol, PPG = polypropylene glycol, NS = normal saline (0.9% NaCl). **(b)** Baricitinib in 50% DMSO/50% PEG-400 that had been incubated at 37°C for 30 days was then tested for inhibitory activity against IL-6-induced Stat3 phosphorylation in human peripheral blood CD4$^+$ T cells. **(c)** B6 mice implanted with baricitinib- or vehicle-loaded osmotic pumps were assayed immediately prior to HSCT (four days post-pump implantation) for IFNγ-induced Stat1 phosphorylation using a whole blood assay.

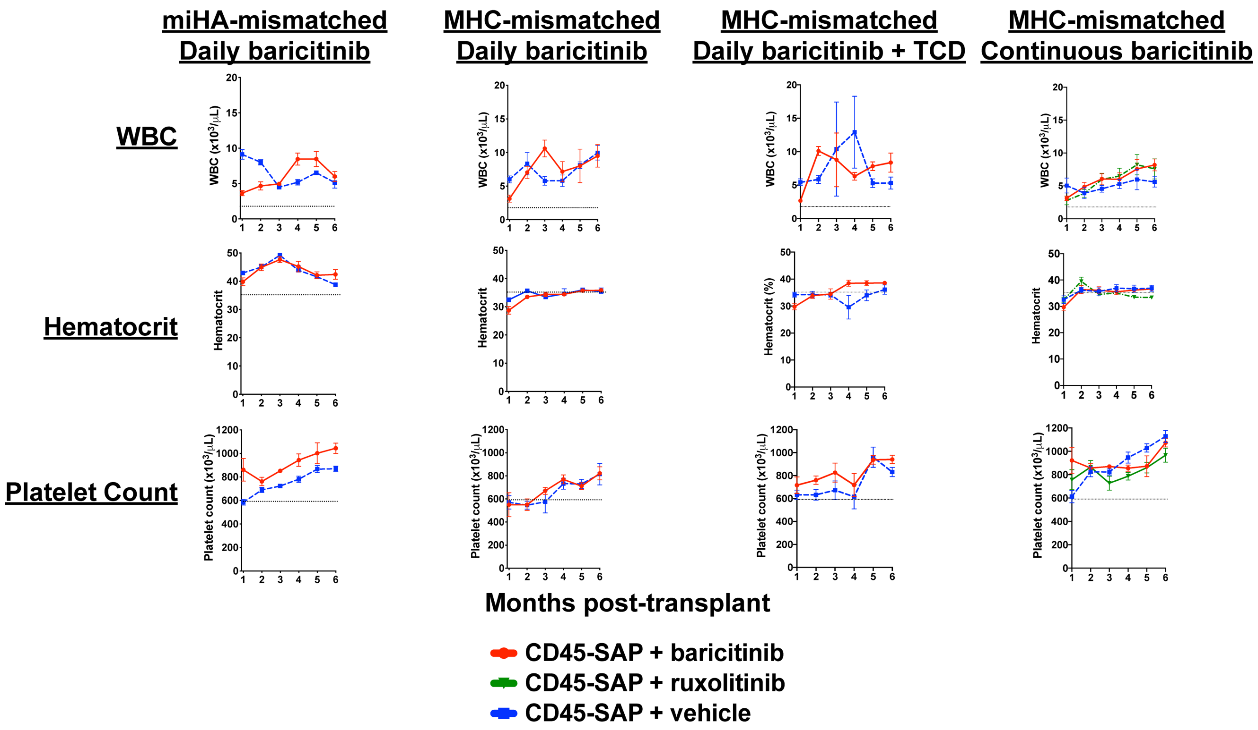


**Supplementary Figure 8. Complete blood counts in allo-HSCT models conditioned with CD45-SAP and JAK1/2 inhibitors.** CBC data for miHA- and MHC-mismatched alloHSCT models dosed daily with baricitinib (first and second columns from left), MHC-mismatched alloHSCT receiving pre-transplant $CD4^+$ and $CD8^+$ TCD plus daily baricitinib (third column), and MHC-mismatched alloHSCT receiving continuously-infused baricitinib via osmotic pump (fourth column) are shown. Dotted lines indicate the lower reference limits for the CBC assays.

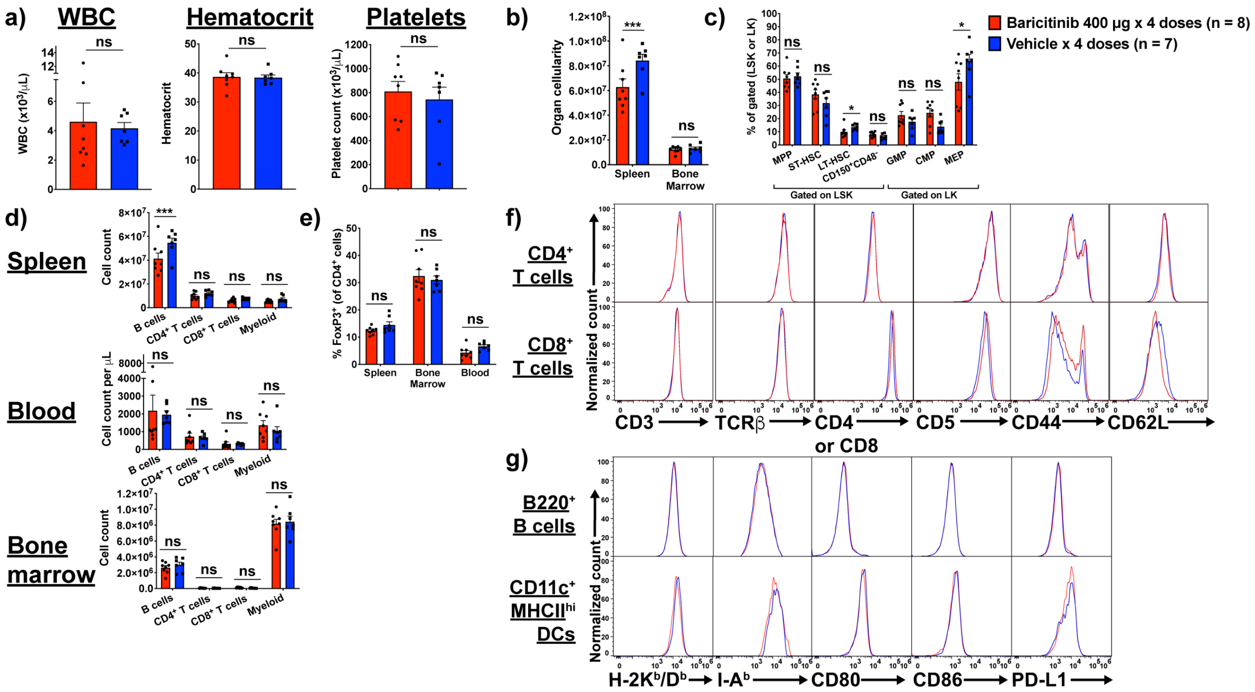


**Supplementary Figure 9. Acute effects of baricitinib on the pre-HSCT recipient environment.** B6 mice were treated daily with baricitinib or vehicle for four days prior to analysis of blood and lymphoid organs. The color scheme in the legend (upper-right) is used throughout the figure. **(a and b)** Complete blood counts (a) and organ cellularity of spleen and bone marrow (b). **(c)** Proportions of HSPC subsets in bone marrow. **(d)** Absolute B, T, and myeloid ($Gr1^+$ and/or $CD11b^+$) cell counts in spleen, blood, and bone marrow. **(e)** Frequencies of $FoxP3^+$ Tregs (relative to total $CD4^+$ T cells) in spleen, blood and bone marrow. **(f and g)** Cell surface phenotyping of splenic T cells (f) and APCs (g). Data points and error bars represent mean ± SEM, with mice pooled across 3 experiments. For statistical comparisons: ns = not significant, * is $p < 0.05$, ** is $p < 0.01$, *** is $p < 0.001$, and **** is $p < 0.0001$.
